# Genetic underpinnings of host manipulation by *Ophiocordyceps* as revealed by comparative transcriptomics

**DOI:** 10.1101/2020.01.03.893917

**Authors:** Ian Will, Biplabendu Das, Thienthanh Trinh, Andreas Brachmann, Robin Ohm, Charissa de Bekker

## Abstract

The ant-infecting *Ophiocordyceps* fungi are globally distributed, host manipulating, specialist parasites that drive aberrant behaviors in infected ants, at a lethal cost to the host. An apparent increase in activity and wandering behaviors precedes a final summiting and biting behavior on to vegetation, positioning the manipulated ant in a site beneficial for fungal growth and transmission. Notably, across *Ophiocordyceps* species and other known host manipulators, the molecular mechanisms underlying behavioral changes remain largely unclear. We explored possible genetic underpinnings of host manipulation by: (*i)* producing a hybrid assembly of the *Ophiocordyceps camponoti-floridani* genome, (*ii*) conducting laboratory infections coupled with RNAseq of both *O. camponoti-floridani* and its host, *Campontous floridanus*, and (*iii*) using these data for a comparative analysis to similar work performed in *Ophiocordyceps kimflemingiae* and *Camponotus castaneus*. We propose differentially expressed genes tied to ant neurobiology, odor response, circadian rhythms, and foraging behavior may be the result of putative fungal effectors such as enterotoxins, aflatrem, and mechanisms disrupting nutrition-sensing or caste-identity pathways.

## Introduction

Transmission from one host to the next is a crucial step in the life cycle of parasites. Certain parasites have evolved to adaptively manipulate the behavior of their animal hosts to aid transmission. Many examples of manipulating parasites and their hosts have been reported across taxa and are active topics of research (Moore 1995, Thomas & al. 2010, Lafferty & KURIS 2012, Moore 2013, Poulin & Maure 2015, de Bekker & al. 2018), with the ant-manipulating *Ophiocordyceps* fungi emerging as a notable model (de Bekker, Merrow, & al. 2014, de Bekker 2019). However, in most parasitic manipulation systems, including *Ophiocordyceps*-ant interactions, the mechanisms by which the parasite dysregulates animal behavior are largely unknown. As such, the study presented here seeks to home in on the major players involved in *Ophiocordyceps* infection and manipulation of carpenter ants by using a comparative transcriptomics framework to identify, compare, and discuss candidate genes underlying manipulation across two different fungus-ant species interactions. The species we compare are *Ophiocordyceps kimflemingiae* and its host *Camponotus castaneus*, on which mechanistic work has previously been performed (de Bekker, Quevillon, & al. 2014, de Bekker & al. 2015, Fredericksen & al. 2017, Loreto & Hughes 2019, Mangold & al. 2019), and *Ophiocordyceps camponoti-floridani* (Araújo & al. 2018) and its host *Camponotus floridanus*. The interactions between the latter have not been investigated in detail yet and we present an initial investigation here.

Ant-manipulating *Ophiocordyceps* infect ants and subsequently modify their behavior to improve parasite transmission at a lethal cost to the host. Infected ants display hyperactivity or enhanced locomotor activity (ELA) (Hughes & al. 2011, de Bekker & al. 2015), deviation from foraging trails (Pontoppidan & al. 2009, Hughes & al. 2011), and a summiting behavior coupled with biting and clinging to attach themselves to vegetative substrates until death (Andersen & al. 2009, Pontoppidan & al. 2009, Hughes & al. 2011, Mongkolsamrit & al. 2012, Chung & al. 2017, Andriolli & al. 2018, Loreto & al. 2018). This final fatal change in behavior is the most tractable readout for manipulation of the host, and this novel behavior provides a growth and transmission site that appears to be adaptive for the fungal parasite (Andersen & al. 2012, Loreto & al. 2014). Bioactive compounds with neuromodulatory and physiology-disrupting effects have been proposed as possible means of dysregulating host behavior (de Bekker, Quevillon, & al. 2014, de Bekker & al. 2015, Kobmoo & al. 2018, de Bekker 2019, Loreto & Hughes 2019), as has tissue destruction and hypercontraction of jaw muscles (Hughes & al. 2011, Fredericksen & al. 2017, Mangold & al. 2019). Moreover, manipulated biting appears to be synchronized in a time of day manner in multiple *Ophiocordyceps*-ant species interactions (Hughes & al. 2011, de Bekker, Merrow, & al. 2014, de Bekker & al. 2015, de Bekker, Will, & al. 2017, de Bekker 2019). This suggests that these parasites also employ mechanisms to modify host behaviors that operate according to daily rhythms and are under control of the hosts’ biological clocks (Hughes & al. 2011, de Bekker, Merrow, & al. 2014, de Bekker & al. 2015, de Bekker, Will, & al. 2017, de Bekker 2019).

Multiple reports indicate that manipulation of ant behavior only occurs in a host-specific manner, with a single species of *Ophiocordyceps* manipulating a single species of ant (Evans & al. 2011, de Bekker, Quevillon, & al. 2014, Araújo & al. 2018, Sakolrak & al. 2018). Part of the mechanisms involved in manipulation of ant behavior might, therefore, be species specific (de Bekker, Ohm, & al. 2017). However, convergently evolved and conserved mechanisms are likely also shared among these specialized *Ophiocordyceps* fungi as they have common evolutionary histories (Araújo & Hughes 2019) and are confronted with similar ecological obstacles (i.e. the modification of ant behavior to attach to elevated transmission sites) (Chetouhi & al. 2015, Loreto & al. 2018). Investigating these shared mechanisms across *Ophiocordyceps* and their host ant species would elucidate some of the common elements involved and help identify candidate genes and compounds that are key to establishing manipulation. Comparative proteomics to understand manipulated phenotypes induced by mermithid worms have demonstrated that such approaches can identify candidate convergent mechanisms of host manipulation across taxa (Herbison & al. 2019). Similarly, here we apply comparative transcriptomics to attempt to reveal candidate genes underscoring manipulation that could offer new insights into ant neurobiology and behavior, novel fungal bioactive compounds, and specific understanding of *Ophiocordyceps* – *Camponotus* interactions. Moreover, such comparative studies may provide evidence and hypotheses for molecular mechanisms driving similar manipulation phenotypes in other systems. Baculoviruses that manipulate the behavior of moth larvae also elicit host ELA, climbing, and eventual death at an elevated position, thereby dispersing viral propagules (Kamita & al. 2005, VAN HOUTE, VAN OERS, & al. 2014, Han & al. 2018). Two primary viral genes have been proposed to be necessary in driving manipulation in this system, an ecdysteroid UDP-glucosyl transferase (*egt*) (Hoover & al. 2011, Han & al. 2015, Ros & al. 2015) and protein tyrosine phosphatase (*ptp*) (Kamita & al. 2005, Katsuma & al. 2012). Another example of a fatal summiting phenotype as a result of parasitic manipulation is induced by the distantly related *Entomophthora muscae* fungi that infect and manipulate flies (Krasnoff & al. 1995, Elya & al. 2018). As *Ophiocordyceps,* Baculovirus, and *Entomophthora* are presented with the similar challenge of inducing a summit disease in insect hosts to improve parasite transmission; quite plausibly parasite mechanisms of manipulation may have convergently evolved.

In the study presented here, we infected *C. floridanus* with *O. camponoti-floridani* and performed RNAseq on both organisms sampled before infection, during manipulated clinging, and after host death. Subsequently, we compared the generated gene expression data to previous transcriptomics work done in *O. kimflemingiae* and *C. castaneus* (de Bekker & al. 2015). Both *Ophiocordyceps* species reside in different clades within the *Ophiocordyceps unilateralis* species complex (Araújo & al. 2018), which are genomically vastly different (de Bekker et al., 2017). With this framework, we transcriptionally compare fungal parasites and ant hosts, highlighting possible shared mechanisms involved in manipulation. To this end, we also report the first annotated genome assembly of *O. camponoti-floridani* using a long-read short-read hybrid approach and transcriptome analyses. We propose candidate fungal genes and possible scenarios by which they may contribute to infection and manipulation of *Camponotus* hosts. Similarly, we highlight host genes that possibly reflect changes in behavior and challenges to physiology due to fungal infection and manipulation. Our findings include changes in genes associated with ant neurobiology, odor detection, and nutritional status, as well as fungal genes related to toxins, insulin-signaling pathways, proteases, and putative effectors similar to those previously reported in Baculovirus. We propose possible scenarios by which these genes may reflect or promote changes in host behavior and physiology as further evidence or new grounds for hypotheses in the field of behavior manipulating parasitism.

## Methods

### Fungal isolation & culture

To sequence the genome and perform infection studies followed by transcriptome sequencing, we isolated and cultured two strains of the fungus *Ophiocordyceps camponoti-floridani*. Strain EC05, used for the de novo genome assembly, was collected from Little Big Econ State Forest in Seminole County, Florida. Strain Arb2 was collected at the University of Central Florida arboretum in Orange County, Florida and used for laboratory infections and RNAseq data. These samples were obtained by permission from the University of Central Florida and the Florida Department of Agriculture and Consumer Services.

Both strains were isolated by surface sterilizing infected *Camponotus floridanus* cadavers in 70% ethanol for 10 sec and aseptically removing ant cuticle with 25 G needles (PrecisionGlide, BD) to extract *O. camponoti-floridani* mycelium. Extracted fungal masses were plunged into a solid medium (7.8 g/L potato dextrose agar [BD], 6 g/L agar [Difco], 100 mg/L Penicillin/Streptomycin [Gibco], and 100 mg/L Kanamycin [Alfa Asear]) and maintained at 28°C for 15 days to screen for possible contaminants and indications of sample viability. We placed viable extractions into liquid culture in T25 tissue flasks (CytoOne, USA Scientific) containing Grace’s Insect Medium (Unsupplemented Powder, Gibco) supplemented with 10% Fetal Bovine Serum (FBS) (Sterile Filtered US Origin, Gibco). Incubation at 28°C and 50 rpm promoted blastospore growth. Once the culture was established, we reduced FBS to 2.5% for secondary cultures.

Both EC05 and Arb2 nuclear 18S ribosomal short subunit (SSU) sequences matched voucher JA-2017c Flx1 (Araújo & al. 2018) with 100% identity, confirming these strains as *O. camponoti-floridani*. We used SSU primers NS1 and NS4 (White & al. 1990), which yielded an approximately 1kb PCR amplicon with a Phusion High Fidelity Polymerase (New England Biolabs) and the following PCR protocol: initial denaturation at 98°C for 30 sec, 30 cycles of 98°C for 10 sec, 49°C for 30 sec, 72°C for 30 s, and final elongation at 72°C for 10 min.

### Whole genome sequencing and assembly

Strain EC05 was used to generate a high-quality draft genome for *O. camponoti-floridani* through a combination of Nanopore long-read and Illumina short-read sequencing. To extract DNA, we disrupted blastospore pellets frozen in liquid nitrogen with a 1600 MiniG tissue homogenizer (SPEX) at 1300 rpm for 30 sec. Samples were processed in 2 mL microcentrifuge tubes (Greiner) containing two steel ball bearings (5/32” type 2B, grade 300, Wheels Manufacturing) and kept frozen throughout disruption. We extracted DNA with 0.9 mL Extraction Buffer (1% SDS [Fisher Scientific], 240 mg/L para-aminosalicyclic acid [ACROS], 484 mg/L Tris/HCl [Fisher Scientific], 365 mg/L NaCl [Fisher Scientific], and 380 mg/L EGTA [MP] at pH 8.5) and 0.9 mL phenol/chloroform (Fisher Scientific). After phase separation, we washed the water phase with chloroform (Alfa Aesar) prior to extracting DNA with isopropanol. Following a 70% ethanol wash and reconstituting in nucleotide-free water (Gibco) we treated the DNA samples with RNase (Thermo Scientific).

A short-read DNA library was prepared with the Nextera DNA Flex Library Prep Kit (Illumina) with an average 390 bp fragment. Indexing for paired-end reads was performed with Nextera i5 and i7 adapters (Nextra Index Kit Index – 1 and 2). Short-read sequences were generated by sequencing 300 bp paired-end reads on an Illumina MiSeq (v 3, Miseq Reagent Kit), resulting in 8 GB of fastq data. Reads were then quality filtered and adapter trimmed using BBduk (Bushnell 2019) as a plugin through Geneious Prime (v 2019.0.3, Biomatters) (trimq = 15, minlength = 75).

To facilitate long-read sequencing, we first size selected genomic DNA for fragments longer than 5 kbp on a Blue Pippin (Sage Science) with 0.75% agarose and a High-Pass protocol. A long-read library was subsequently generated using the SQK-LSK109 Ligation Sequencing Kit (Oxford Nanopore) according to manufacturer’s protocols. Sequencing on a PromethION (R9 flowcell, Oxford Nanopore) generated 105 GB (estimated 180x coverage) of Nanopore sequence data. Sequencing reads were base called with Albacore (v 2.2.5, Oxford Nanopore) and adapters were trimmed with Porechop (Wick 2018). We assembled the initial long-read genome using Canu (v 1.7.1, genomeSize = 45m, default settings) (Koren & al. 2017). An overestimation of the genome size allowed us to generate an assembly with good coverage despite the presence of bacterial contaminants (see below). This initial Canu long-read assembly was polished using raw Nanopore read data through Nanopolish (v 0.10.2) (Loman & al. 2015), followed by Illumina reads (120x coverage) with three iterations of Pilon (Walker & al. 2014) (v 1.23, --fix all) to produce a hybrid assembly. We identified a putative mitochondrion contig by testing for circular sequence structure with Circlator (Hunt & al. 2015), MUMmer (Kurtz & al. 2004), and Canu (Koren & al. 2017).

The assembly contained bacterial contaminant contigs that we removed. We identified contaminant contigs by their: (*i*) low read coverage aligned with Minimap2 (Li 2018) using all EC05 reads (Nanopore average coverage: 553x of *O. camponoti-floridani* genome contigs and 58x of contaminant contigs, and, Illumina: 195x of genome contigs and 15x of contaminant contigs); (*ii*) low RNA coverage with HISAT2 (Kim & al. 2015) mapping of Arb2 RNAseq control culture samples (62x of *O. camponoti-floridani* genome contigs and 0.14x of contaminant contigs); and (*iii*) high mapping to known bacterial genomes, *Cohnella* sp. 18JY8-7 (GenBank CP033433.1), *Delftia acidovorans* isolate ANG1 (GenBank CP019171.1), and *Stenotophomonas maltophilia* strain ISMMS2 (GenBank CP011305.1) (0.08% overlap between *O. camponoti-floridani* genome contigs and these bacteria genomes, and 28.96% overlap of contaminant contigs with these bacteria genomes).

### Genome annotation

We predicted 7455 genes in the EC05 *O. camponoti-floridani* genome using Augustus (v 3.0.2) trained with BRAKER1 (v 1.1.8) and intron hints from Arb2 transcripts (Stanke & al. 2008, Hoff & al. 2016). Protein domains predicted by PFAM (v 32) (Finn & al. 2014) were used to identify associated GO terms (Ashburner & al. 2000, Hunter & al. 2009). Protease predictions were made with the MEROPS database and a BLASTp E-value cutoff of 1e-5 (Rawlings & al. 2014). We used TMHMM (v 2.0c) to annotate transmembrane domains (Krogh & al. 2001). Secretion signals were identified with SignalP (v 4.1) (ALMAGRO Armenteros & al. 2019). We predicted small secreted proteins (SSPs) when genes were shorter than 300 amino acids, carried a SignalP secretion signal, and did not have a transmembrane domain outside the first 40 amino acids. We identified genes and clusters predicted to be involved in secondary metabolism using a pipeline based on SMURF (Khaldi & al. 2010, de Bekker & al. 2015), with parameter d = 3000bp and parameter y = 6. Transcription factors were identified based on the presence of a PFAM domain with DNA-binding properties using PFAM mappings from (Park & al. 2008). For BLAST annotations, we used BLASTp (v 2.7.1) against the NCBI nr database to gather up to 25 hits with E-value ≤ 1e-3. For the final annotation, we passed these hits to the Blast Description Annotator of Blast2GO with default settings (Conesa & al. 2005). In addition to searches that returned no results, we considered descriptions starting with “hypothetical protein” or “predicted protein” to lack BLAST annotations. BLASTp searches of the predicted proteins of *O. camponoti-floridani* against the Pathogen-Host Interaction (PHI) database (Urban & al. 2017), mitochondrial proteins, fungal secondary metabolite cluster proteins, and the *O. kimflemingiae* genome were conducted using Geneious (v 2019.0.3, Biomatters) with E-value ≤ 1e-3 and bit-score ≥ 50.

We supplemented the published BLAST genome annotations of the latest version of the *C. floridanus* genome (v 7.5) (Shields & al. 2018) with PFAM and GO annotations using the InterPro database (Finn & al. 2017) through Blast2GO (Conesa & al. 2005). For these additional annotations, we used the longest transcript variant per gene. To allow for comparison of ant RNAseq results of our study to DE BEKKER & al. (2015), we bridged the current *C. floridanus* assembly to the earlier version (v 1.0) (Bonasio & al. 2010) used by DE BEKKER & al. (2015) through BLASTp homology searches with Geneious (v 2019.0.3, Biomatters), taking the top hit after an E-value ≤ 1e-3 and bit-score ≥ 50 cutoff.

### Ant collection & husbandry

Ant infections and behavioral observations were done using a wild colony of *C. floridanus*. This colony was collected from the University of Central Florida arboretum in February 2018 and housed in the laboratory. The colony consisted of several hundred individuals including minors, majors, and brood. In order to acclimate the ants and entrain their biological clocks to laboratory conditions, we first subjected the colony to two days of constant light and constant 25°C temperature in a climate controlled room. Following this clock “reset” we gave the colony three days of 12 hr – 12 hr light-dark cycles at 25°C to entrain ants to light as a circadian zeitgeber (LD1212, lights begin at zeitgeber time ZT 0). During acclimation, the colony housed in a 9.5 L plastic container (42 cm long x 29 cm wide) lined with talcum powder (Fisherbrand) and containing aluminum foil wrapped test-tubes (50 mL, Fisherbrand) with moist cotton to serve as darkened, humid nest spaces. Ants fed *ad libitum* on 15% sucrose solution, autoclaved crickets, and water.

### Ant infections

For laboratory infections, we selected minor caste ants from the colony and housed them in two identical containers in each of two climate-controlled incubators. Incubator A (MIR-154) was programmable for light and temperature. Incubator B (I36VL, Percival) was programmable for light, temperature, and relative humidity (RH). Incubator A ran a program with LD1212 and 28°C during the light phase and 20°C during the dark phase. Incubator B maintained humidity at 70% RH, LD1212, and 28°C to 20°C temperature. The light phase of Incubator B included a 4hr increasing ramp step (ZT 0 dark and 20C transitioning to ZT 4 peak light and 28°C), a 4hr peak light and temperature hold until ZT 8, and 4hr decreasing ramp step until ZT 12 (peak light and 28°C to dark and 20°C). Light, temperature, and humidity for both incubators were verified with a HOBO data logger (model U12, Onset) (Supplementary fig.1a-b). The incubators were not significantly different for survival of infected ants (p = 0.072, log-rank test). Therefore, we chose to consolidate all samples from these incubators for survival and RNAseq analysis.

Each container (33 cm x 22 cm) was lined with talcum and had a thin layer of playground sand on the bottom that we routinely moistened during observations to maintain an elevated humidity inside the ant enclosure. On one end, containers held a 50 mL Falcon tube (Corning) with moist cotton wrapped in aluminum foil and *ad libitum* 15% sucrose and water. On the opposite end, we placed two thin 12 cm high wooden sticks draped with locally collected “Spanish moss” (*Tillandsia usneodies*) and a single “air plant” (*Tillandsia spp.*) (Supplementary fig.1c). These plants are common natural substrates for manipulated ants to bite and cling to at local field sites.

We painted ants to distinguish treatment groups (POSCA paint pens, Uni) one to three days in advance of infection by fungal injection (de Bekker, Quevillon, & al. 2014). We injected ants without anesthesia using aspirator tubes attached to glass capillary needles (10 µL borosilicate capillary tubes, Fisherbrand), pulled using a PC-100 Narishige instrument. Needle placement for injection was on the ventral side of the thorax, sliding under the prosternum. We removed sugar and water from infected ants the night before injection to ease the procedure. We timed injections to begin at ZT 0 and not last more than 3.5 hr. Ants that survived the first 3 hr post-injection were placed into the experiment. Control ants were not injected. Sham treatment ants were injected with 1 µL of Graces-2.5% FBS. Infected ants were injected with 1 µL of 3.5 x 10^7^ blastospores/mL in Graces-2.5% FBS, harvested during log-phase growth (OD_660nm_ = 0.984, approximately 1.8 x 10^7^ cells/mL based on estimates with *Saccharomyces cerevisiae*). Blastospores were harvested immediately preceding injection, washed twice in deionized water, and re-suspended in Graces-2.5%FBS.

Incubator A contained 18 control, 13 sham, and 33 infected ants. Incubator B contained, 12 control, 13 sham, and 30 infected ants. After 14 days post injection (dpi), we observed aggressive patrolling and cleanup of dead and dying ants. Therefore, we chose to separate infected ants from non-infected groups to reduce the chances of interference with cadavers or the progress of manipulation. Control ants and sham ants were removed from the experiment boxes and rehoused in similar containers directly next to their original box for the remainder of the experiment.

### Observations of infection progression and sample collection

We made daily observations for manipulated ant phenotypes and survival at ZT 0, 2, 4, 6, 8, and 23 with sporadic opportunistic surveys for manipulated ants. We additionally began observations at ZT 20 starting 18 dpi. We considered ants to be manipulated when they displayed clasping or biting onto any substrate. Individuals that ceased to move nor responded to agitation by air puffs were considered dead. Live manipulated ants collected for RNAseq were recorded as dead for survival analysis. We analyzed survival data using the R package survival (Therneau 2015) and visualized curves with survminer (Kassambara & al. 2019).

Upon visible behavioral manipulation, we froze whole-ant samples for RNAseq directly in liquid nitrogen. We sampled healthy live control ants at ZT 21, which corresponds to the time of observed manipulation in our study. Healthy controls, rather than sham-injected ants, were collected to better match the previous study on *O. kimflemigniae* and *C. castaneus*, which we reference for comparative transcriptomics (de Bekker & al. 2015). Upon flash freezing, we stored ants in pre-chilled 2 mL microcentrifuge tubes (USA Scientific) at −80°C until RNA extraction. In total, we analyzed 13 ants for RNAseq: live manipulated n = 5, dead manipulated n = 5, healthy control n =3. To obtain fungal control samples of strain Arb2 (n = 3), blastospore cultures were harvested at ZT 21 after a constant light and 28°C synchronization treatment for two days, followed by an entrainment period for five days at LD1212 and 28°C to 20°C. During this time, light and temperature of culture conditions were validated with a HOBO (Supplementary fig.1b). Fungal control cultures were grown to a late-log phase (OD_660nm_ = 1.7) before harvesting by pelleting 1 mL of culture per sample and snap-freezing in liquid nitrogen. Any collections made during subjective dark were done under red-light (730 nm wavelength).

### RNAseq data generation and analysis

All frozen samples for RNAseq were disrupted in the same manner as fungal genomic DNA samples (see above) prior to RNA isolation. For ant samples, we first decapitated frozen cadavers in petri dishes chilled with liquid nitrogen and then proceeded to frozen tissue disruption using individual heads. We extracted RNA with a RNAqueous Micro kit (Life Technologies) according to the manufacturer’s protocol, without DNase treatment. We isolated mRNA with poly-A magnetic beads (NEB) from 500 ng total RNA for each sample. Subsequently, we converted purified mRNA to 300 bp fragment DNA libraries with the Ultra II Directional kit (NEB) and indexed them for multiplexing (NEB).

All libraries were sequenced on an Illumina HiSeq as 100 bp single-end reads, resulting in 27M to 56M reads for each sample. We trimmed reads using BBduk (Bushnell 2019) as a plugin through Geneious Prime (v 2019.0.3, Biomatters) to remove adapters and for quality (qtrim = rl, trimq = 10, minlength = 25). Our choice for a Q10 quality trim and minimum 25 bp length of RNAseq reads yielded a sufficient number and quality of reads while reducing risk of introducing biases from read processing (Williams & al. 2016).

For mixed transcriptome libraries (infected ants with host and parasite reads), we conservatively separated transcript sequences by first discarding all reads from the mixed sample that mapped to one organism’s genome before proceeding to analyze the other organism’s transcriptome (Fig.1). That is, we would map to the host genome and then align the unmapped reads to the parasite genome, and *vice versa*. This method removes reads that map ambiguously to both the host and parasite from analysis. However, we estimate this to be only ≤ 0.04% of reads based on these organism’s transcriptomes in control conditions (Fig.1). All transcript mapping steps were done with HISAT2 (Kim & al. 2015).

**Figure 1.**
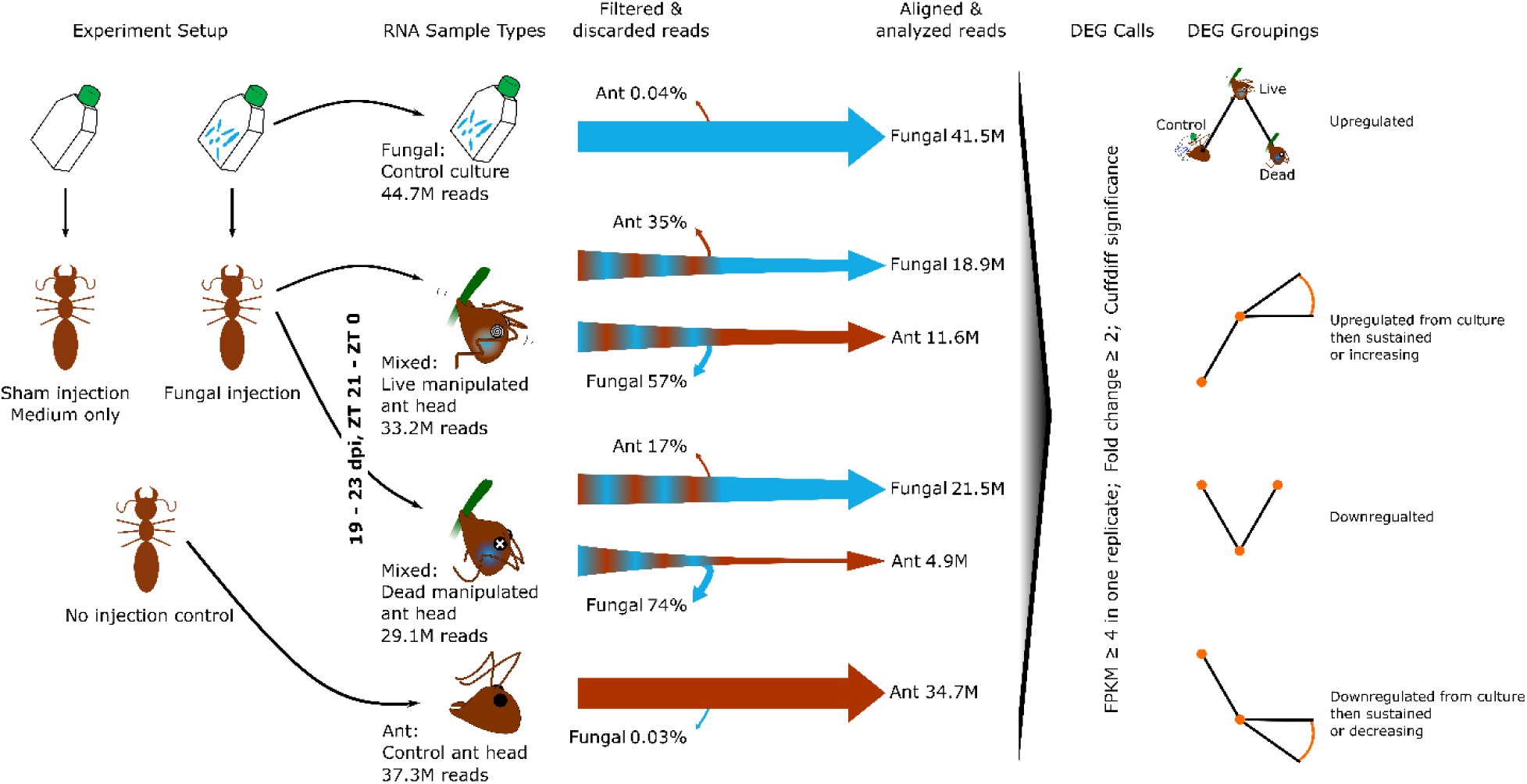
RNAseq experimental overview. Three ant treatment groups, fungal injection, sham injection, and no injection were generated at the start of the experiment. Sham injection treatments were used in survivorship analysis only. Fungal injections, ant controls (no injection), and fungal controls (blastospore culture) produced samples used for RNA extraction. Manipulated ants contained mixed RNA samples of both the parasite and host. We observed and collected manipulation samples 19 – 23 dpi and between ZT 21 to ZT 0. Control ants and fungal culture samples were collected at ZT 21. To minimize possible bias from read counts of transcripts from mixed samples that map to either organism, we first filtered reads through either the host or parasite genome before aligning remaining reads to the other genome for analysis. DEGs were identified with Cuffdiff and expression cutoffs we imposed for comparability to De Bekker *& al. (2015)*. DEGs were then grouped by expression pattern over the course of the different sample points. All RNA samples taken from ants were extracted from whole heads. Read counts are the mean value per sample type.

We normalized and analyzed whole transcriptome gene expression levels with Cuffdiff with default settings (Trapnell & al. 2012). In addition to using Cuffdiff significance calls (q ≤ 0.05, test = OK, significant = yes) to identify differentially expressed genes (DEGs) between sample groups, to consider genes biologically relevant to our analyses we required genes to have an expression level of ≥ 4 FPKM in one replicate, and a minimum of two-fold change between sample types. This methodology allows us to have the most comparability to published RNAseq data from *O. kimflemingiae* and *C. castaneus* (de Bekker & al. 2015)

We used unsupervised principal component analyses (PCA) to describe the variation among control, live manipulated, and dead manipulated samples using R (R CORE TEAM 2014, RSTUDIO TEAM 2015). We ranked genes within a principal component (PC) by loading values to investigate the top 20 that explain the most variation within a PC. All gene FPKM transcription values were first *log*2(*X* + 1) transformed for every gene with at least one sample replicate with FPKM ≥ 4.

Using a weighted gene coexpression network analysis (WGCNA) we produced modules of coexpressed genes and associated them with control, live manipulated, and dead manipulated ants. Although our sample size is below an ideal replicate number, we applied this analysis for a coarse evaluation of possible gene modules associated with manipulation. Data were filtered (FPKM ≥ 4) and *log*2(*X* + 1) transformed before analysis. We processed all samples together in the R package WGCNA (v 1.67) (Langfelder & Horvath 2008). We applied a signed-hybrid network type, a soft power threshold equal to nine (fungus) or 12 (ant), minimum module size of 30, and default settings. Our categorical trait data were entered as either 0 (sample was not that type) or 1 (sample was that type) for control, live manipulation, and dead manipulated. For correlation of ant and fungal modules, eigengene values for ant modules were calculated with the moduleEigengenes function of the package and used as trait data for fungal module correlations. R package WGCNA was also used to generate a sample dendrogram to asses clustering of biological replicates (function hclust, with method = “average”).

We performed enrichment analyses on gene sets identified by PCA, WGCNA modules, and DEG groupings by using a hypergeometric test with Benjamini-Hochberg correction to correct for multiple testing (minimum number of genes with annotation term = 5, corrected p-value ≤ 0.05).

## Results

### Manipulated behavior of *Camponotus floridanus* after laboratory infections

We set out to identify parasite and host genes involved in manipulated biting and clinging behavior observed in *Ophiocordyceps-*infected *Camponotus* ants. As such, we aimed to perform comparative transcriptomics between two *Ophiocordyceps* species and their respective *Camponotus* hosts. To this end, we infected *C. floridanus* with *O. camponoti-floridani* to compare gene expression levels in this host-parasite interaction with those published for *C. castaneus* and *O. kimflemingiae* (de Bekker & al. 2015).

All manipulated *C. floridanus* ants (n = 11) clung to plants with their legs, with two individuals additionally biting the plant. This lab-infected manipulated phenotype appears to well-approximate wild manipulations (Fig.2). Live manipulated ants commonly displayed subtle tremors, feeble clasping motions, and low responsiveness to puffs of air. Additionally, if these ants fell from their manipulation perches, they continued clasping motions but were otherwise unable to right themselves or move (n = 2). In addition to generating manipulations, overall survival was significantly different based on treatment (p = 0.00059, log-rank test), with the 95% confidence interval of infected ants lower and not overlapping with sham treated or control ants by 21 dpi (Fig.2).

**Figure 2.**
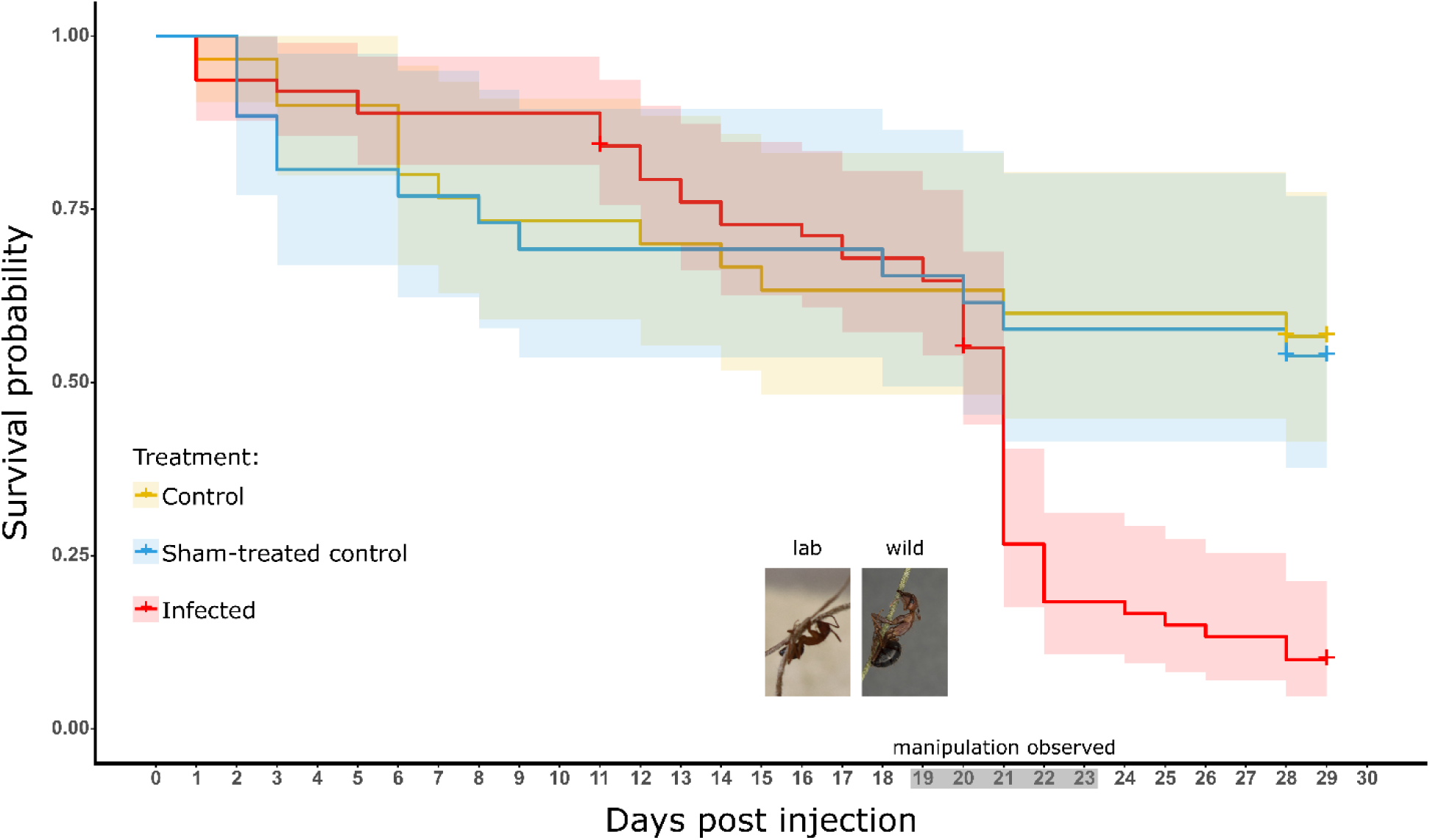
Survival curves of ants not injected (yellow, control n = 30), injected with media-only (blue, sham-treated control n = 26), or injected with the *O. camponoti-floridani* (red, infected n = 63). Shaded regions around survival data indicate 95% confidence intervals. Treatment had a significant effect on survival (p = 0.00059, log-rank test). Manipulations were observed only in infected ants, 19 to 23 dpi (gray axis shading). Crosses indicate censorship events when ants were removed from the experiment and no longer contributed to analysis. The photo inserts show a live lab infected ant (left) and a wild manipulated cadaver (right).

Manipulated clinging and biting behavior displayed by *O. camponoti-floridani* infected *C. floridanus* ants occurred within a stereotypic dpi-window with apparent time of day synchronization. Infected ants displayed manipulated behavior starting 19 dpi, with the last manipulation at 23 dpi (Fig.2). All observed manipulations occurred pre-dawn from at least as early as ZT 20 to as late as ZT 23 (i.e., 4 hr – 1 hr before lights-on). In cases where we captured the onset of manipulated clinging and allowed the ant to progress to death, the time between manipulation and death was 0.5 hr to 2 hr (n = 3, of six dead manipulated ants). Under the laboratory conditions used in infection studies with *O. kimflemingiae*, manipulations of *C. castaneus* occurred 16 through 24 dpi and shortly after subjective dawn (ZT 3), which was the first daily observation period of that study (de Bekker & al. 2015). Infected *C. castaneus* usually died at least 5 hr after manipulation. Such stereotypic patterns have also been reported for other ant-manipulating *Ophiocordyceps*, both in the laboratory (Sakolrak & al. 2018) and in the wild (Hughes & al. 2011).

Our preliminary field observations have found live manipulated *C. floridanus* one to three hours after solar noon (n = 4). This is out of phase from our laboratory observations for this species, as well as those made for *O. kimflemingiae*-infected ants (de Bekker & al. 2015). Differences in abiotic factors, such as light, temperature, and humidity, across labs and between experimental and field observations, are plausible influences that could have led to these shifts in timing (Hughes & al. 2011, Andriolli & al. 2018, Cardoso Neto & al. 2019). However, conclusive observations to reveal which cues may regulate the onset of manipulation still need to be accumulated. Thus, rather than selecting a comparable ZT at which to sample infected *C. floridanus* for RNAseq, we sampled infected ants based on comparable phenotypes, immediately upon observing manipulated clinging behavior or death after manipulation. Sampling according to phenotype instead of daily timing should result in more comparable gene expression profiles across the two species-interaction studies and, therefore, reveal the key genetic components they have in common.

### De novo hybrid assembly of the Ophiocordyceps camponoti-floridani genome

To reliably align and separate sequencing reads of mixed samples and determine the relative abundance of both host and parasite transcripts requires high-quality reference genomes representing both organisms (Fig.1). A recently updated genome of *C. floridanus* is publicly available (Shields & al. 2018). However, prior to this study, a genome of *O. camponoti-floridani* has not yet been produced. To generate a high-quality *O. camponoti-floridani* genome we combined Nanopore long-reads and Illumina short-reads data in a hybrid assembly.

After polishing and contaminant removal steps, our *de novo* hybrid assembly contained 13 contigs encompassing 30.5 Mbp, with a N50 of 3.8 Mbp and 53x Nanopore coverage (Tab.1). Pezizomycotina telomeric repeats, TTAGGG (Podlevsky & al. 2008), are present on eight of the contigs, three of which are bounded on both ends by repeats and therefore should represent whole chromosomes of 5.6 Mbp, 3.8 Mbp, and 2.5 Mbp in length. The assembly appeared nearly complete with 99.7% of pezizomycotina benchmarking universal single-copy orthologs (BUSCOs) (v 3, using OrthoDB v. 9) (SIMÃO & al. 2015) (Tab.1) (Accession # in progress). The larger size of this assembly (30.5 Mbp) compared to *O. kimflemingiae* (23.9 Mbp) (de Bekker, Ohm, & al. 2017) did not result in an increased number of predicted genes (7455 and 8629, respectively) (Tab.1). This size difference could reflect true variation in the composition of these *Ophiocordyceps* genomes. Alternatively, the *O. camponoti-floridani* hybrid assembly has been able to better parse out repeat regions, describing their length and frequency more accurately, than the short-read assembly on which the *O. kimflemingiae* genome is based. Using hybrid assembly techniques to generate high quality, large contigs generally improves representation of genomic architecture. This can be important for understanding repeat regions, duplications, and gene linkage relationships such as secondary metabolite clusters. Indeed, *O. camponoti-floridani* metabolite cluster 8 (see secondary metabolite section below) seemed to comprise two distinct clusters in *O. kimflemingiae* (de Bekker, Ohm, & al. 2017) as a single homologous unit. This finding could again be a real difference between the two species, but likely indicates that the long-read data assembled into a more complete *Ophiocordyceps* genome.

One of the 13 contigs represents a putative mitochondrion of 272,497 bp. We identified this mitochondrial contig by *(i)* its high read coverage (735x) compared to the genome average (53x), *(ii)* its low GC content (27%) compared to the total assembly (48%), *(iii)* the presence of homologs to known mitochondrial proteins on this contig but nowhere else in the genome (BLASTp of ATP6, COB, COX1, and NAD1 of *Aspergillus niger*) (Joardar & al. 2012), and *(iv)* its circular sequence structure. The circular structure was detected through three approaches: Circlator (Hunt & al. 2015), self-alignment with MUMmer (Kurtz & al. 2004), and Canu (Koren & al. 2017). Circlator and Canu did not detect circular contigs elsewhere in the assembly. Circularizing based on MUMmer overlaps reduced this contig from 384,563 bp to 272,497 bp.

Genome annotations identified 7455 gene models (Tab.1). Most genes received functional annotations based on PFAM domains (73%) or BLAST descriptions (86%). We also identified 801 putatively secreted proteins containing a SignalP domain (11%) and 271 small secreted proteins (SSPs, 3.6%). Only 19% of the SSPs carried known PFAM domains and only 37% returned BLAST descriptions. With many SSPs lacking clear functional annotations, these SSPs may contain a pool of novel bioactive compounds secreted by *O. camponoti-floridani*.

### RNAseq identifies differentially expressed genes associated with manipulation

To discover candidate fungal and ant genes that underpin the manipulated behavior of *O. camponoti-floridani*-infected *C. floridanus* hosts, we sequenced the transcriptomes of samples obtained before, during, and after manipulation. Ant heads collected during and after manipulation contained mixed transcriptomes of both host and parasite (Fig.1). The average aligned reads for each analyzed fungal and ant transcriptome fell between 11.6M and 41.5M, except for the ant transcriptome of the dead manipulated samples (i.e. after manipulation), which resulted in only 4.9M aligned reads (Fig.1). For both fungal culture and healthy ant head control samples, 93% of the RNAseq reads aligned to their respective reference genomes (Shields & al. 2018). We aligned 57% and 74% of reads obtained from live manipulated and dead manipulated ant heads to the *O. camponoti-floridani* genome, respectively. In contrast, these samples only resulted in 35% and 17% of reads that aligned to the *C. floridanus* genome (Fig.1) (Shields & al. 2018). These findings suggest that the fungus has colonized the ant head by the time of manipulation and rapidly destroys host tissue for its own growth as the host dies, as has been observed in other *Ophiocordyceps* – ant interactions (Hughes & al. 2011, de Bekker & al. 2015).

To validate differential expression analysis between our biological sample groups, we first determined if the variation between replicates within these groups was smaller than the variation between them. Unsupervised dendrograms based on replicate ant and fungal gene expression profiles indeed clustered biological replicates together (Supplementary fig.2). However, the fungal profile of live manipulation sample 4 (L4) was placed ambiguously relative to live and dead manipulated samples (Supplementary fig.2b). Regardless, we did not exclude sample L4 as an outlier, as we are not confident about the confines of typical disease progression and gene expression during this time point. As these samples were selected on observed behavioral phenotype, the transcriptional state of genes essential for manipulation is plausibly shared despite differences in other genes. We included all samples to identify differentially expressed genes correlated to control (fungal culture and healthy ants), live manipulation, and dead manipulation samples. Using Cuffdiff (Trapnell & al. 2012) significance (q ≤ 0.05, test = OK, significant = yes), FPKM expression minimums (FPKM ≥ 4), and fold change thresholds (≥ 2-fold), we found that 1431 ant and 2977 fungal genes were differentially expressed between at least one pair of sample conditions (control, live manipulation, or dead manipulated). To identify genes that are plausibly involved in *Ophiocordyceps* manipulation of ant behavior, we performed various complimentary gene expression analyses for both fungal and ant genes, which are detailed below.

### Principal component analyses identify host changes linked to manipulation

Through PCAs we identified the genes that contributed the most to the transcriptome variations between our biological groups. An unsupervised PCA of all normalized ant transcriptome data distinguished host gene transcription prior to infection (control), during manipulation (live manipulation), and after (dead manipulated) (Fig.3a). Principal Component 1 explained 47% of the variation between host transcriptional profiles over progression of the infection from healthy to manipulated to dead ants. To identify the major contributors to this PC describing the change in hosts from healthy, to manipulated, to dead, we ranked and plotted ant genes by their PC1 loading values Fig.3b, Supplementary tab.1). The top 20 genes of PC1 included genes putatively related to odor detection (pheromone-binding protein Gp-9-like, and a pheromone-binding protein (PBP)/general odorant-binding protein (GOBP) family PFAM domain-containing gene), nutrition and energy balance (alpha-amylase 1, alpha-amylase A-like, and an apolipophorin-III precursor PFAM domain-containing gene), muscle tissue (myogenesis-regulating glycosidase-like, and muscle actin), and cell growth (CREG1). These genes were also found to be differentially expressed between sample types and are further highlighted in the DEG section below.

**Figure 3.**
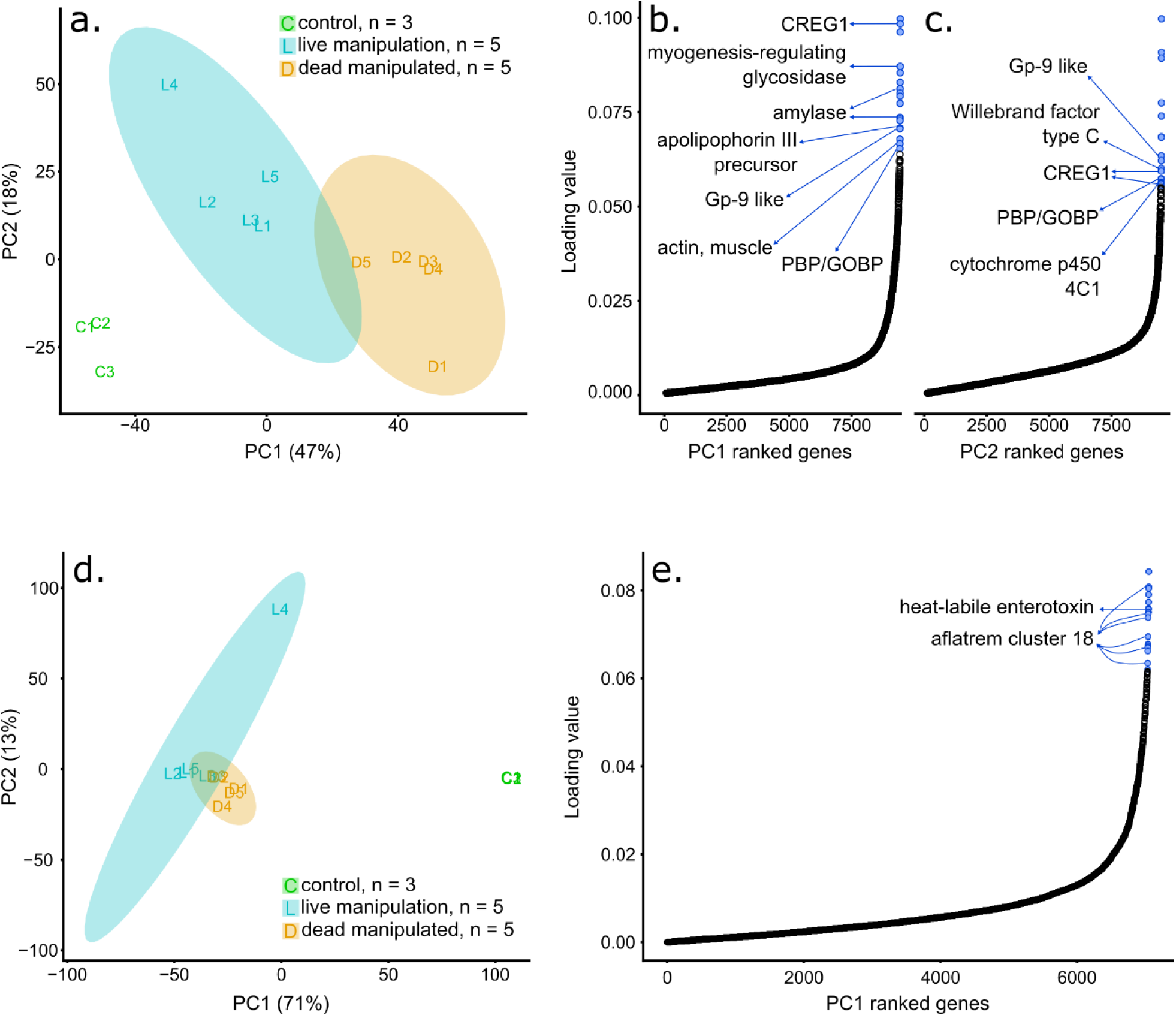
Principal component analyses and plotting of principal component loading values from normalized ant (*a*, *b*, *c*) and fungal (*d*, *e*) gene expression values. PCA plots (*a*, *d*) show gene expression from controls (green), during live manipulation (blue), and after host death (orange). Shaded regions indicate 95% confidence ellipses; control samples do not have sufficient sample size to produce an ellipse. *(a.)* Replicates of host gene expression largely cluster and vary across PC1 as the state of the host progresses from healthy control ants to live manipulated hosts to dead manipulated hosts killed by the fungus. PC2 primarily describes the variation between healthy controls and live manipulated ants, but all biological groups vary along this axis. *(b, c)* All ant genes ranked by loading values in PC1 (*b*) or PC2 (*c*). Most genes have low loading values, however a relatively high-loading value subset contribute the most to PC1. Of these high value genes, the top 20 include genes with marked increases in loading values that may play key roles during infection and manipulation (blue). *(d.)* Gene expression of the fungal parasite interacting with the ant host is clearly distinguished from that of fungal culture control samples by PC1. *(e.)* All fungal genes ranked by loading values in PC1. The top 20 fungal PC1 genes are also highlighted (blue).

PC2 explained 18% of the variation and rather highlighted the difference between control and live manipulation than control and dead manipulated hosts. The top 20 PC2 genes, which largely describe the variation between healthy control ants and live manipulated hosts, shared some transcriptomics signals with PC1 (i.e., PBP/GOBP domain carrying genes and CREG1 genes). However, the PC2 top 20 also included putative DEGs involved in insect immunity (defensin and a von Willebrand factor type C domain-containing gene) and insect starvation response-mediated by juvenile hormone (JH) (cytochrome P450 4C1-like) (Fig.3c, Supplementary tab.2).

### Principal component analyses identify fungal effectors produced during infection

A PCA of all normalized fungal transcriptome data displayed a large separation between transcription profiles prior to infection (control) and after (live or dead manipulated) (Fig.3d). Fungal gene expression profiles from live and dead manipulated ants were less different from each other, as indicated by the partial overlap of the 95% confidence ellipses of these biological groups (Fig.3d). Fungal PC1 explained 71% of the transcriptional variation between growth in control cultures and inside ants’ heads Fig.3d). The second principal component (PC2, 13%) largely described the difference between replicate L4, and other fungal samples, which our dendrogram also indicated (Supplementary fig.2). As such, major elements of PC1 likely indicate genes linked to infection, manipulation, and killing of the host.

Upon ranking and plotting all fungal genes by their PC1 loading values, we selected the top 20 for inspection (Fig.3d, Supplementary tab.3). For every gene in this set, peak transcript FPKM values occurred during live manipulation in both *O. camponoti-floridani* and *O. kimflemingiae* homologs (de Bekker & al. 2015). Although significantly higher than gene expression in culture, their differential expression relative to dead host samples was not always significant. This set of 20 genes contained multiple genes of interest identified in secondary metabolite clusters and as DEGs. These genes, discussed in more detail in the sections below, included multiple members of a putative aflatrem biosynthesis pathway (cluster 18) and a putative enterotoxin that was found to be extremely highly up-regulated in both *O. camponoti-floridani* and *O. kimflemingiae*.

### Weighted gene coexpression network analysis correlates parasite gene expression during manipulation to host gene networks

In addition to performing PCAs, we analyzed both ant and fungal RNAseq data using a WGCNA to describe coexpressed gene modules correlated with control, live manipulated and dead manipulated samples. For ant gene coexpression patterns, the WGCNA generated 22 modules, which we named A1 – A22. Four of the modules, A4, A5, A6, and A10, were significantly positively correlated with live manipulation (p ≤ 0.05, Fisher’s asymptotic test on Pearson correlation values) and either had a negative or no significant correlation with control and dead samples (Supplementary fig.3). Four additional modules, A14, A15, A17, and A18, were significantly negatively correlated to live manipulation. Of these modules, A17 and A18 were also positively correlated to healthy control ants. Their downregulation during manipulation could, thus, represent the reduced expression of gene networks associated with a healthy state, which are either suppressed by the fungus or as a host-response to infection. Taken together, the WGCNA identified eight ant gene modules with significant correlations to the time of live manipulation that highlight transcriptional responses to fungal infection and manipulation (Supplementary data files). The WGCNA for fungal gene coexpression patterns generated 13 modules, F1 – F13, and correlated them to the three possible sample types – control culture, live manipulation, and death after manipulation. Three modules, F1, F2, and F4, were significantly positively correlated with live manipulation. F1 and F2 were additionally negatively correlated with fungal growth in control culture (Supplementary fig.4). Subsequently, we performed an additional WGCNA with all 13 fungal modules against the eight ant modules that had the most meaningful correlation patterns to host condition to describe possible behavioral changes and responses to infection. We used these eight ant modules as a new set of trait data (i.e., eigengenes) to correlate our fungal modules to (Fig.4). Using this strategy rather than separately associating fungal and ant gene networks to the broader categories of our biological groups, we aimed to make a more detailed connection between fungal gene expression and the corresponding transcriptional changes in the host.

**Figure 4.**
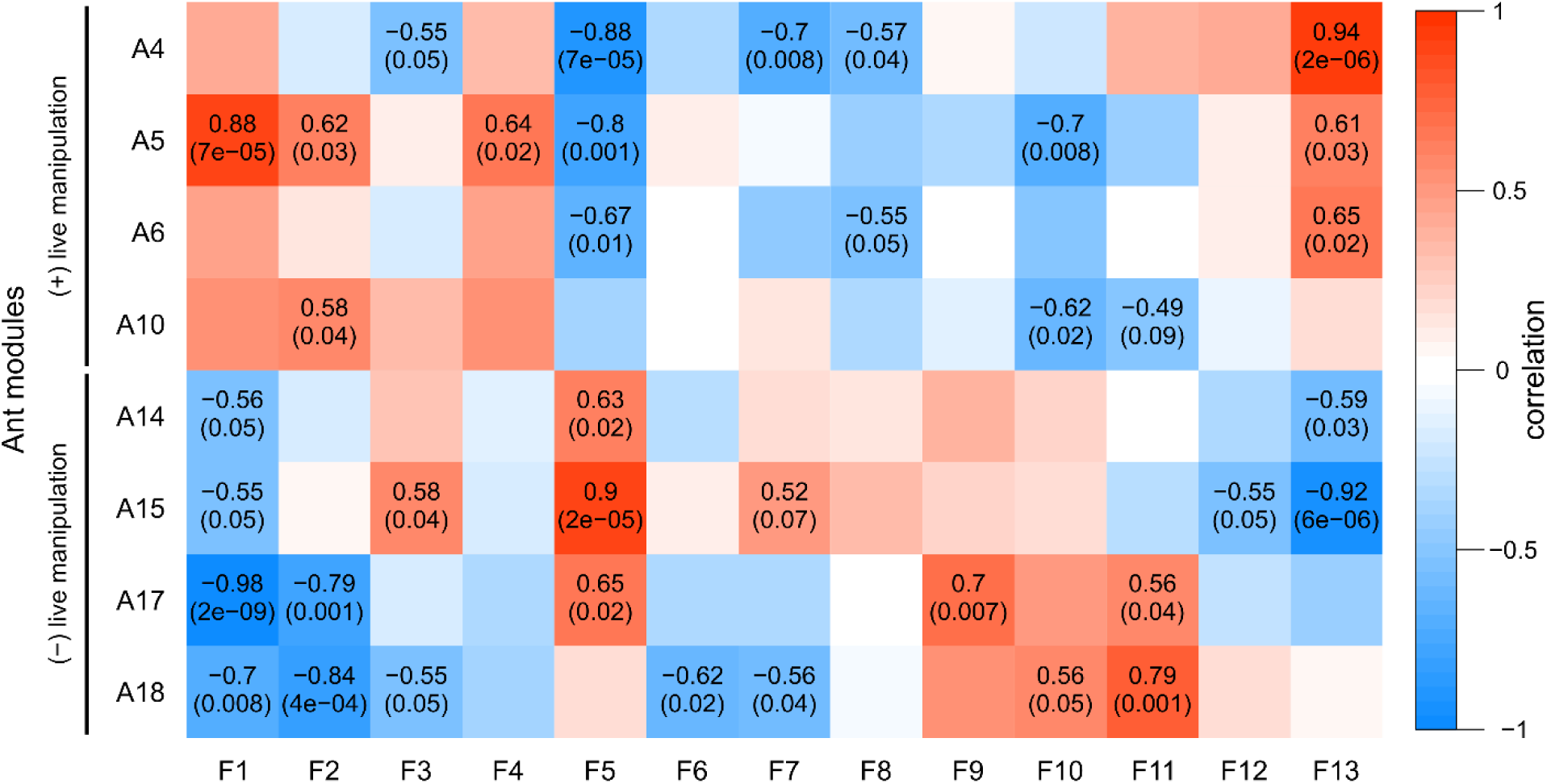
WGCNA of fungal gene expression (columns) showing correlation (color and top number in each cell) and p-value (in parentheses) of gene modules to selected ant modules that are either positively or negatively correlated to samples during live manipulation (rows). Correlation and p-values are shown only for significant module to module correlations. Modules F1, F2, and F13 are enriched for secretion signals or transmembrane domains and therefore may contain extracellular fungal effectors. Modules A14 and A15 have PFAM domain overrepresentations suggesting a role in neuronal function and development.

To investigate the broad functions of our identified gene modules, we performed enrichment analyses of annotated PFAM domains and GO terms present in those modules. In ant module A4, genes annotated to be involved in odorant detection (i.e., PBP/GOBP PFAM domain) and with encoded proteasome activity were overrepresented (Supplementary data files). These gene functions were also highlighted by the ant PCA above. Reduced detection of odor cues and related social interactions have been hypothesized to play a role in the early stages of manipulation preceding biting by carpenter ants infected with *Ophiocordyceps* (de Bekker & al. 2015). In module A5, gene regulatory functions were enriched as DNA binding, transcription factor, and DNA replication related annotations were overrepresented (Supplementary data files). Similarly, modules A6 and A10 showed enrichment of gene regulation domains and terms, however, with fewer such terms than module A5 (Supplementary data files).

Of the ant modules negatively correlated to live manipulation, A14 and A15 had a 7 transmembrane receptor (rhodopsin family) PFAM enrichment related to light sensing and cellular signaling (Supplementary tab.4). This suggests a loss in light sensitivity as the mechanism promoting light-seeking behavior, which has previously been hypothesized to occur in manipulated ants prior to biting to assure light levels that promote fungal growth and transmission (Andriolli & al. 2018). Module A15 additionally contained an overrepresentation of genes with a 7 transmembrane domain (sweet-taste receptor) related to glutamate or GABA receptors, and, neurotransmitter gated ion channels (i.e., genes carried BLAST annotations for acetylcholine, glycine, and glutamate receptors) (Supplementary data files). Moreover, both modules were predominantly enriched for GO terms and PFAM domains related to transmembrane transport, ion regulation, and cell signaling activity with multiple immunoglobulin domain enrichments (Supplementary data files) – although the underlying genes for these enrichments were generally not DEGs. The immunoglobulin domain overrepresentations contained a variety of genes putatively encoding cell-surface binding proteins related to neuronal development, maintenance, and activity, such as IgLON family proteins with additional light sensing or circadian, olfaction, and memory related functions (Supplementary tab.5). Taken together, neurotransmitter related genes were notably present in modules A14 and A15. This indicates that these gene networks are closely tied to the ant’s neurobiology and that changes in the relative expression of these networks may be linked to manipulation. The negative correlations of modules A14 and A15 to the time of live manipulation suggest the dysregulation of responsiveness to light and neurotransmitters related to behavior and muscle activity as mechanisms underlying the manipulated behaviors observed.

Fungal modules F1, F2, and F13 appeared to be major contributors to fungal effects on the ant host as these modules were positively correlated to activation of manipulation-associated ant modules and negatively correlated to ant modules deactivated during manipulation (Fig.4). Putatively secreted genes (i.e., SignalP and SSP annotations) were overrepresented in F1 and F2, as were transmembrane domains (TMHMM) in modules F1 and F13. These enrichments suggest that these fungal modules are involved in extracellular interactions with the host. Oxidation-reduction related GO terms and transcription factors were also overrepresented in module F2. Oxidation-reduction terms are a hallmark of parasite-host interactions and were also overwhelmingly found for *O. kimflemingiae* – *C. castaneus* interactions (de Bekker & al. 2015). Although fungal modules F1 and F13 did not harbor any other annotation enrichments, they indicated an increase of putatively secreted effectors and virulence related activity in *O. camponoti-floridani* in correspondence to the changing expression of ant gene networks correlated with manipulation. Notably, F1 and F13 negatively correlated with ant modules A14 and A15, which in turn appear to be associated with neuron function. Therefore, modules F1 and F13 possibly contain extracellular fungal effectors that dysregulate neuron function and health.

It is worth noting that due to sample size constraints (n =13), the module results of our WGCNA may contain some artifacts. We, therefore, limited our analysis to general module content and correlation, rather than including more detailed approaches to identify hub genes. The networks produced by the WGCNA contained many genes with high intramodular connectivity, module membership, and trait correlation with no clearly informative cutoff to define hub genes by. For example, a membership cutoff of > 0.8 still retained 170 of 274 genes in fungal module F1, and 677 of 891 in F2. As such, we choose not to propose hub genes for these modules. However, including the WGCNA as one of the multiple analytical approaches to describe gene coexpression (i.e., through PCA, WGCNA, and DEG groupings) enabled us to home in on the more robust candidate manipulation genes that surface across different analyses. Our WGCNA indeed shares major patterns with the other analyses we performed as it linked increased compound secretion to manipulation and produced modules that contained plausible infection and manipulation-related DEGs upregulated during infection (see below). Full WGCNA enrichments to add in supp file, BIP, need A1-3, 7-9, 11, 19-22

### Differentially expressed ant genes during infection and manipulation

To consider genome-wide enrichment patterns in ant gene expression in relation to infection, manipulation and host death, we divided our RNAseq data into subsets representing different interpretations of gene function that may underlie observed effects or be host responses to fungal activity (Fig.1). Genes that were significantly upregulated from control ants to live manipulation and then downregulated once the ant died may indicate specific responses to infection and active behavioral manipulation by the parasite (120 genes, Supplementary data files). We also analyzed genes that were upregulated during live manipulation from control and then had sustained or increased transcription until the ant died (88 genes, Supplementary data files). Being upregulated, these genes plausibly play a role during manipulation but may also be more generally associated with infection or host death. Similarly, we considered differentially expressed gene sets that were downregulated at live manipulation relative to both healthy ants and dead hosts (6 genes, Supplementary data files), and those holding or dropping further in dead hosts (529 genes, Supplementary data files). As we collected our samples for the dead manipulated time point under frequent observation, these ants should reflect a manipulated ant transcriptome just as the host dies. As significant transcriptional effects would likely lag behind the moment of presumed death, we expect to have captured the state of a moribund ant as it dies. To propose host changes that could be underlying the manipulated behaviors in *C. floridanus*, we closely investigated the functional annotations of these DEG sets and compared them to those previously found for manipulated *C. castaneus*.

#### Host gene expression patterns related to starvation and tissue destruction

Over the course of infection by *Ophiocordyceps*, ant hosts are expected to lose energy stores to the parasite. The expression of genes related to nutritional balance will likely reflect this change in physiology. Moreover, starvation often induces hyperactivity in animals exhibited by increased levels of locomotion behavior (Yang & al. 2015), which in some cases could be adaptive for the host (Hite & al. 2019) or reflect parasitic manipulation. Indeed, we found DEGs putatively related to primary metabolism and nutrition in *Ophiocordyceps*-infected ants, which we frequently observe to display elevated bouts of locomotion prior to death. In starved *Drosophila* flies, lipase 3, an enzyme involved in fat metabolism, was found to be up-regulated (Zinke & al. 1999). Similarly, we identified an upregulated lipase 3 gene (2-fold increase from control to live manipulation) in ants sampled during manipulation. However, an additional four lipase 3 genes and an overrepresentation of the GO term “lipid metabolic process” among the downregulated subset of genes suggested that overall lipid metabolism might be diminishing in manipulated ants. Indeed, two putative apolipophorin genes (apolipophorin and an apolipophorin III precursor domain containing gene) were also significantly downregulated over the course of infection and manipulation in both *C. floridani* and *C. castaneus* (between 2-fold to 9-fold decrease from control to live manipulation) (de Bekker & al. 2015). Apoliophorin III has immune functions that operate in a trade-off manner with metabolism of high fat diets in insects (Adamo & al. 2010).

In addition to signs of a diminishing lipid metabolism, we found other possible hallmarks of starvation that could result in increased locomotion. One of them is cytochrome p450 4C1, which has been implicated in starvation responses in cockroaches and responds to JH treatments in females (Lu & al. 1999). Although in cockroaches, starvation led to the upregulation of this gene, we found thirty putative cytochrome p450 4C1 genes downregulated from control to live manipulation in *C. floridanus*, with three homologs also significantly downregulated in *C. castaneus* (de Bekker & al. 2015). An alpha amylase A-like gene, which is involved in the degradation of complex sugars (Da Lage 2018), was also downregulated over the course of infection (286-fold decreases from control to death in *C. floridanus* and 12-fold in *C. castaneus*). This indicates a reduction of starch metabolism in the likely starving host. Also tied to nutritional state, we observed a putative *insulin-1* gene downregulated in both species of ant hosts (3-fold and 6-fold decrease from control to live manipulation in *C. floridanus* and *C. castaneus*, respectively). As we note changes in gene expression not in line with reports in other insects (i.e., lipase 3, apolipophorin III, and cytochrome P450 4C1), this could mean that the ants’ energy reserves have been fully depleted at our nearly-terminal time points of sampling. Alternatively, genetic starvation mechanisms are different in ants compared to flies and cockroaches, or the fungal parasite has disrupted the ants’ typical starvation responses. Possibly, changes in nutritional status also underlie behavior changes in the host, such as ELA, that may be adaptive for the parasite. This scenario is discussed in more detail in the differentially expressed fungal gene section below.

Besides the modification of nutritional physiology, tissue destruction also occurs during infection. Notably, the ants’ muscles are affected, leading to muscle atrophy and hypercontraction (Hughes & al. 2011, Fredericksen & al. 2017, Mangold & al. 2019). In line with this aspect of the disease, a putative sarcalumenin gene was downregulated during live manipulation in both *C. floridanus* and *C. castaneus* (3-fold and 8-fold decrease from control to live manipulation, respectively) (de Bekker & al. 2015). Sarcalumenin has been shown to interact with calcium and regulate muscle excitation and fatigue in mammalian systems (Zhao & al. 2005, O’Connell & al. 2008). Additionally, an essential component of myosin motor proteins, a myosin light chain alkali (Yamashita & al. 2000), was downregulated in both species of ant hosts during live manipulation (4-fold and 2-fold decrease from control to live manipulation in *C. floridanus* and *C. castaneus*, respectively). Similarly, the expression levels of three ant myogenesis-regulating glycosidase genes significantly decreased from healthy control ants, to live manipulated, and to dead manipulated ants, as well as a gene annotated as “actin, muscle” (10-fold decrease from control to live manipulation). However, its homolog in *C. castaneus*, followed the opposite expression pattern. Taken together, the expression levels in both host species show hallmarks of muscle tissue destruction and dysregulation in ant heads of individuals that were manipulated to display biting and clinging behaviors. These genes could be part of the fungus-induced muscle mechanisms that drive these behaviors.

#### Shifts in caste identity and circadian rhythms that control suites of behavior

Ants within the same colony are genetically highly similar. Despite this, ants in different castes show vastly different caste-related behaviors. For instance, the nursing caste stays inside the nest to groom and feed the brood while the foraging caste leaves the nest at set times to forage for food. One could therefore predict that changes in host behavior during *Ophiocordyceps* infection result from changes in the expression of genes related to caste behaviors. Juvenile hormone (JH) is known to shape the formation of caste identity during development in eusocial hymenoptera and continues to correlate with caste behaviors into adulthood (Ament & al. 2008, Chandra & al. 2018, Opachaloemphan & al. 2018). In our transcriptome dataset, we found two JH activating genes (juvenile hormone acid O-methyltransferases) and two deactivating genes (juvenile hormone epoxide hydrolases) that were significantly downregulated in live manipulated ants compared to healthy controls (Shinoda & Itoyama 2003, Zhang & al. 2005). A similar gene expression pattern was found for a homologous JH epoxide hydrolase in *C. castaneus* (de Bekker & al. 2015). Regulating cell growth and apoptosis, CREG1 has also been proposed to modulate the development of worker-reproductive caste status in a JH responsive manner (Barchuk & al. 2007, Beckstead & al. 2007, Wurm & al. 2010). All three genes putatively encoding for CREG1 in the *C. floridanus* genome were downregulated over the course of infection, two of which had similarly regulated homologs in *C. castaneus* (de Bekker & al. 2015). Taken together, *Ophiocordyceps*-induced changes in ant behavior could be partially due to the dysregulation of JH levels.

Ecdysteroid kinases are enzymes that inactivate ecdysteroids that interact with JH and can indirectly influence insect development and behavior via this route as well (Libbrecht & al. 2013). Modification of ecdysteroids has been implicated in the behavioral manipulation of moth larvae by baculovirus to assure that they remain in an elevated position (Hoover & al. 2011, Han & al. 2015, Ros & al. 2015). We detected an enrichment of genes containing ecdysteroid kinase domains in the subset of ant genes that were downregulated from control to live manipulation and remained lowly expressed or decreased further into host death. The modification of ecdysteroids could, thus, also be involved in *Ophiocordyceps* manipulation of ant behavior to induce climbing behavior, and perhaps be a more general mechanism underlying summiting in parasite-manipulated insects.

Ant circadian rhythms have been put forward as a possible host aspect for *Ophiocordyceps* to dysregulate as part of manipulation (de Bekker & al. 2015, de Bekker, Will, & al. 2017, de Bekker 2019). Healthy ants display daily regimented foraging behaviors controlled by the molecular clock. These behaviors are seemingly disrupted and replaced by manipulated climbing, biting, and clinging behaviors that in turn take place in a synchronized manner (Hughes et al., 2011, de Bekker et al., 2015). As controls and manipulated ants were time-matched in both this study and DE BEKKER & al. (2015), differential expression of clock-related genes are likely due to infection by *Ophiocordyceps* and not an artifact of time of day during sampling. In line with the circadian clock hypothesis, we found that core clock gene, circadian locomotor output cycles protein kaput (*clk*) (Darlington & al. 1998) was significantly downregulated from healthy control to live manipulated ants in both *C. camponoti-floridani* (10-fold decrease) and *C. castaneus* (3-fold decrease) (de Bekker & al. 2015). Additionally, a gene putatively encoding the clock controlled Takeout (TO) protein showed a similar gene expression pattern, again both in *C. floridanus* and *C. castaneus* (3-fold and 4-fold decreases from control to live manipulation, respectively) (de Bekker & al. 2015). Takeout is a JH interacting protein involved in insect foraging behaviors, which has been proposed as a possible target for parasitic disruption of insect locomotor activity and behavior (VAN HOUTE & al. 2013). We have found more evidence for this hypothesis here and further discuss possible fungal effectors dysregulating *to* in the fungal DEG section below.

#### Dysregulation of odor detection and sensory related genes

Odor detection is at the basis of social organization and behavior in ants, mediated by multiple odorant receptors and odorant binding proteins (OBPs). Pheromone binding proteins (PBPs) are a subset of OBPs specialized in binding pheromones and as such play a vital role in an individual’s response to external stimuli (VAN DEN BERG & ZIEGELBERGER 1991, Chang & al. 2015). We found 16 such odor receptor and binding protein genes to be differentially expressed in *C. floridanus* (Supplementary data files); 13 of which were differentially expressed between controls and live manipulation, six being downregulated and seven upregulated.

One of the odorant receptors, putatively encoding an odorant receptor coreceptor (ORCO), is highly conserved in insects and has a central role in odor detection (Jones & al. 2005). Dysregulation of *orco* in ants has been linked to changes in overall sensitivity to odorants, and affects behavior such as time spent outside the nest, ability to detect prey, and aggression towards conspecifics (Yan & al. 2017). One of two putative *orco* genes in *C. floridanus* was significantly upregulated in the ant during live manipulation compared to control (3-fold increase). However, its homolog in *C. castaneus* was significantly downregulated (4-fold decrease) (de Bekker & al. 2015). Two additional putative odorant receptor genes in *C. floridanus* were also differentially expressed from control to live manipulation (i.e., upregulated *or1* and downregulated *or4*-like). Homologs of these genes have been proposed to encode for pheromone receptors in moths (Grosse-Wilde & al. 2011, Wicher & al. 2017). The differential expression of these three odor receptors suggest that manipulated *Camponotus* ants suffer from dysregulated odorant detection.

Among the 11 differentially expressed PBP and general-OBP (GOBP) domain containing genes, four were annotated to putatively encode pheromone binding protein Gp9. Variation in *Gp9* influences colony dynamics and behavior by regulating queen number in colonies of fire ants (Ross 1997, Ross & Keller 1998, Krieger & Ross 2002, Gotzek & al. 2007, Gotzek & Ross 2007). Three of these *Gp9-*like genes were significantly upregulated from control to live manipulation in *C. floridanus*, while the fourth was down-regulated. Although the expression profiles of *C. castaneus* homologs did not match in this case, relatable patterns of odorant receptor and OBP dysregulation were found in manipulated *C. castaneus* (de Bekker & al. 2015). As such, *Ophiocordyceps* infected individuals may generally be unable to properly communicate with nestmates and recognize organizational signals. This dysregulation could be facilitating the wandering behaviors we observe in infected individuals. If infected ants are thereby pulled away from the colony and more commonly positioned in suitable transmission sites, communication disruption through odor reception malfunction could be a process that is adaptive to the parasite.

Other genes possibly associated with odor communication were also downregulated from control to live manipulation in *C. floridanus*. One of these genes was a putative *sensory neuron membrane protein 1* (17-fold decrease), which is involved in the detection of lipid-derived pheromones in pheromone-sensing antennal neurons (Pregitzer & al. 2014). We additionally detected two putatively encoding acyl-CoA Delta(11) desaturases (16-and 4-fold downregulated), which are key enzymes in the synthesis of pheromones in moths (Choi & al. 2002) and may speculatively play a role in chemical communication of ants as well. A homolog to the 16-fold downregulated acyl-CoA Delta(11) desaturase was also reduced in expression in *C. castaneus* (1.8-fold decrease) but does not meet our DEG threshold requirements (de Bekker & al. 2015).

We also detected the upregulation of a putative calpain D gene in the ant during live manipulation relative to both control and dead manipulated samples (5-fold increase from control to live manipulation). Its homolog was similarly upregulated in manipulated *C. castaneus* (6-fold increase) (de Bekker & al. 2015). Calpain D plays a role in the development of optic lobe neurons and mutant *Drosophilia* suffer subsequent defects in locomotor behaviors (Fischbach & Heisenberg 1981, Delaney & al. 1991). In addition to odor sensing, light sensing and response have been proposed as possible aspects of host manipulation by *Ophiocordyceps* (Hughes & al. 2011, Andriolli & al. 2018). Changed levels in Calpain D expression might be a sign thereof.

#### Dysregulation of neurotransmitter signaling

Dysregulation of neurotransmitter and neuronal-activity modulating compounds are a plausible parasite strategy to manipulate host behavior. We identified a suite of ant neuron regulating and neurotransmitter receptor genes in the WGCNA modules that were negatively correlated to samples collected at live manipulation (see above). Closely inspecting DEGs, we also identified putative ant neuromodulatory compounds that were differentially expressed over the course of infection.

Kynurenic acid is an anticonvulsant and neuroprotective neuroinhibitor that when dysregulated, has been implicated in mammalian neurodegenerative disease, changes in activity levels, and reduced motor coordination (P. Yu & al. 2004, Yu & al. 2006). A putative kynurenine/alpha-aminoadipate aminotransferase, which promotes the synthesis of kynurenic acid, was upregulated in the ant during live manipulation (3-fold increase from control to live manipulation) as was the homologous gene in *C. castaneus* (33-fold increase) (de Bekker & al. 2015). Additionally, DE BEKKER & al. (2015) found that a fungal kynurenine formamidase was upregulated during live manipulation of *C. castaneus*. This enzyme drives the synthesis of kynurenine, the precursor of kynurenic acid and other compounds. Notably, *O. kimflemingiae* also appeared to promote neuroprotection of the *C. castaneus* brain during manipulation by the secretion of the neuroprotectant ergothionine (Loreto & Hughes 2019).

Biogenic monoamines have neuromodulatory roles in insects, and changes in their activity and synthesis may underlie manipulated phenotypes in ants. Octopamine functions as a neurotransmitter in insects, acting through G-protein coupled receptors (GPCRs). Octopamine has been found to modulate learning and memory, foraging behavior, olfactory decision making, aggression, and social interactions (reviewed in Roeder 2005). Additionally, as discussed above, octopamine has been linked to starvation-induced elevation of locomotion activity (Yang & al. 2015). These are all processes that may be affected in ants manipulated by *Ophiocordyceps* (Hughes & al. 2011, de Bekker & al. 2015). Consistent with this scenario, we identified octopamine receptors that were differentially expressed between live manipulated and control ants. In *C. floridanus*, a putative octopamine receptor beta-2R was downregulated during manipulation (2-fold decrease from control to live manipulation). while beta-3R, was found to be upregulated in *C. castaneus* (6-fold increase from control to live manipulation) (de Bekker & al. 2015). This suggests that octopamine responsiveness is dysregulated *in* Ophiocordyceps-infected ants.

Dopamine, another biogenic monoamine, also functions as a neurotransmitter in insects and regulates motor neuron activity, locomotion behavior, and biting behavior, among other processes, sometimes in a clock-controlled fashion (Cooper & Neckameyer 1999, Ceriani & al. 2002, Szczuka & al. 2013). Tyrosine 3-monooxygenase drives the rate-limiting step in dopamine synthesis (Daubner & al. 2011). We found a gene putatively encoding for this enzyme to be significantly upregulated during manipulation in *C. floridanus* (4-fold increase from control to live manipulation), as was the homologous gene in *C. castaneus* (3-fold increase) (de Bekker & al. 2015). In addition, a putative DOPA decarboxylase, which catalyzes the final step of dopamine synthesis (Daubner & al. 2011) was significantly upregulated in *C. floridanus* during manipulation (3-fold from control to live manipulation). Changes in dopamine levels may also be implicated in immune function, as a precursor to melanin, which is a component of ant immunity (Ratzka & al. 2011). Changes in the sensitivity to and production of biogenic monoamines in ants correlated to manipulated biting suggest that these neurotransmitters play a role in producing the observed behaviors.

In addition to neuroprotective agents and biogenic monoamines, we identified two more differentially expressed genes that could be involved in aberrant neuronal functioning in manipulated individuals. Both *C. floridanus* and *C. castaneus* exhibited reduced expression of a putative *dimmed-*like transcription factor in live manipulated ants compared to the healthy controls (6-fold and 9-fold decrease, respectively) (de Bekker & al. 2015). The downregulation of *dimmed* resulted in the dysregulation of neuropeptide secretion and insulin/insulin-like growth factor (IGF) signaling (IIS)-responsive neuronal maintenance in *Drosophila* (Hamanaka & al. 2010, Luo & al. 2013, Liu & al. 2016). A putative *bubblegum* gene was also downregulated in *C. floridanus* during manipulation (5-fold decrease from control to live manipulation) as was the homolog in *C. castaneus* (4-fold decrease) (de Bekker & al. 2015). Bubblegum, which is an acyl-CoA synthetase that esterifies fatty acids, plays a role in healthy neuron function. In *Drosophila*, *bubblegum* mutants displayed neurodegeneration, retinal degeneration, and reduced locomotor activity (Min & Benzer 1999, Sivachenko & al. 2016).

### Differentially expressed putative fungal effector genes

As for the ant gene expression data, we divided the fungal data into subsets representing different interpretations of gene function in relation to manipulation (Fig.1). Fungal genes that were significantly upregulated from culture to live manipulation and then downregulated once the host died likely play a role in infection and/or behavioral manipulation (307 genes, Supplementary data files). For enrichment analysis, we further narrowed this set of upregulated genes to the top 50^th^ percentile of genes with the largest downregulation in dead hosts (168 genes, Supplementary data files). Genes in this 50^th^ percentile are tightly regulated relative to the manipulation event and, therefore, possibly the most manipulation specific genes in our dataset. We also considered genes that were upregulated during live manipulation from culture and then had sustained or increased transcription in the dead host (1088 genes, Supplementary data files). Being upregulated, these genes plausibly play a role in infection or manipulation. They may also play a role in fungal activities associated with host death, such as killing the host and consuming dead host tissues. Similarly, we considered differentially expressed gene sets that were downregulated at live manipulation relative to both culture and dead hosts (61 genes, Supplementary data files), a 50^th^ percentile strongly down subset (33 genes, Supplementary data files), and downregulated from culture and holding or dropping further in dead hosts (867 genes, Supplementary data files). These gene sets could either indicate genes not important for manipulation, or the reduced transcription of inhibitors.

Of the 307 fungal genes that significantly peaked during live manipulation, compared to fungal culture controls and dead manipulated ant samples, 90 were found to have homologs with a similar expression pattern in *O. kimflemingiae* (de Bekker & al. 2015) (Supplementary data files). This is likely an underestimation of genes that would share this pattern, as the *O. camponoti-floridani* – *C. floridanus* interactions spanned a much shorter time between manipulation and death than the *O. kimflemingiae* – *C. castaneus* interactions (de Bekker & al. 2015). Therefore, genes in the current experiment had less time for a significant difference in transcript levels to accumulate between live manipulation and dead manipulated samples. Eleven of the 61 *O. camponoti-floridani* genes that were downregulated during manipulation had homologs with the same pattern of transcripts in *O. kimflemingiae*. Proportionally, these two species have a greater overlap in genes upregulated during manipulation (29%) than those downregulated (18%). This suggests that upregulation during manipulation contains proportionally more genes with conserved function and fitness constraints, i.e. involvement in infection and manipulation.

We also compared DEGs upregulated and downregulated during manipulation using the Pathogen-Host Interaction (PHI) database (Urban & al. 2017). We counted PHI descriptions that contained any annotation other than “unaffected_pathogenicity” as genes putatively involved in virulence and manipulation. Upregulated genes comprised 57 hits with pathogenicity annotations, while only 22 were present in the downregulated gene set. In both cases, numerous gene products without PHI annotation result were present (201 upregulated, 32 downregulated). Of these genes absent in the PHI database, 54 genes in the upregulated set putatively encoded secreted proteins (i.e., SignalP annotation), whereas only 4 such genes were present in the downregulated group. These genes, that lack PHI database annotation but are part of the secretome, could contain novel undescribed fungal effectors that are relevant to *Ophiocordyceps* – *Camponotus* interactions.

Over 25% of the entire fungal secretome was upregulated from culture to living manipulated ants (Supplementary tab.6). However, the function of many of these proteins remains elusive. Fungi in live manipulated ants upregulated 239 genes with SignalP secretion signals relative to culture (129 had a PFAM annotation), 85 of which represent SSPs (only 19 had a PFAM annotation) (Supplementary tab.6). DE BEKKER & al. (2015) identified 195 genes putatively encoding secreted proteins upregulated during live manipulation by *O. kimflemingiae*, which represents an upregulation of 22% of its putative secretome. The increased activation of the *Ophiocordyceps* secretome during manipulation by both species suggests a critical role for secreted compounds in modifying host behavior. This is in line with microscopy evidence demonstrating that fungal cells do not grow invasively into the ant’s brain (Hughes et al., 2011; Fredericksen, 2017), but rather likely manipulate behavior peripherally by secreting neuroactive compounds. However, the specific functions of many putatively secreted proteins, upregulated during manipulation, have yet to be determined. Indeed, 74% (23 of 31) of upregulated SSPs lacked both PFAM and BLAST annotations.

Enrichment analyses of DEG sets yielded candidate classes of genes with plausible links to infection and manipulation. No annotation terms were enriched among the downregulated genes. However, we identified multiple genes related to cell cycle and reproduction, which could indicate that changes in fungal growth activity take place during manipulation as well. In fungal genes upregulated during live manipulation relative to culture, or both culture and dead manipulated samples, we uncovered putative effectors of infection, manipulation, and host killing. These effectors are discussed in more detail below.

#### Ophiocordyceps upregulate ADP-ribosylating toxins and ribotoxin during manipulation

Heat-labile enterotoxins are AB ADP-ribosylating agents that have been described for pathogens such as *Escherichia coli*, *Cordyceps bassiana*, and *Metarhizium robertsii* (reviewed in Lin & al. 2010, Mannino & al. 2019). These toxins are formed by an enzymatic A-subunit and five structural cell-binding B-subunits. Heat-labile enterotoxins in *E. coli* are known to modify GTP-binding proteins and interfere with G protein-coupled receptors and subsequent intra-cellular signaling through increased cyclic AMP levels. This process eventually leads to cell dysfunction and apoptosis (reviewed in Lin & al. 2010, Mangmool & Kurose 2011).

The *O. camponoti-floridani* genome contains 35 predicted heat-labile enterotoxin genes based on PFAM annotation (PF01375|Enterotoxin_a). Of these genes, 30 also carry SignalP secretion domains, 10 of which were upregulated from culture to live manipulation, and then six were sharply downregulated in the dead host. A seventh putative enterotoxin, that lacked the characteristic PFAM domain but did have a heat-labile enterotoxin blast description, was similarly expressed. As such, we found heat-labile enterotoxin alpha chain domains to be enriched among upregulated genes during manipulation (Fig.5). This group of genes also resulted in the enrichment of the GO terms “multi-organism process,” “interspecies interactions between organisms,” “toxin activity,” and “pathogenesis” (Supplementary data files). We additionally found enterotoxins to be present in the manipulation associated fungal WGCNA modules F1 (n = 3) and F2 (n = 8, including the most upregulated enterotoxin, discussed below).

**Figure 5.**
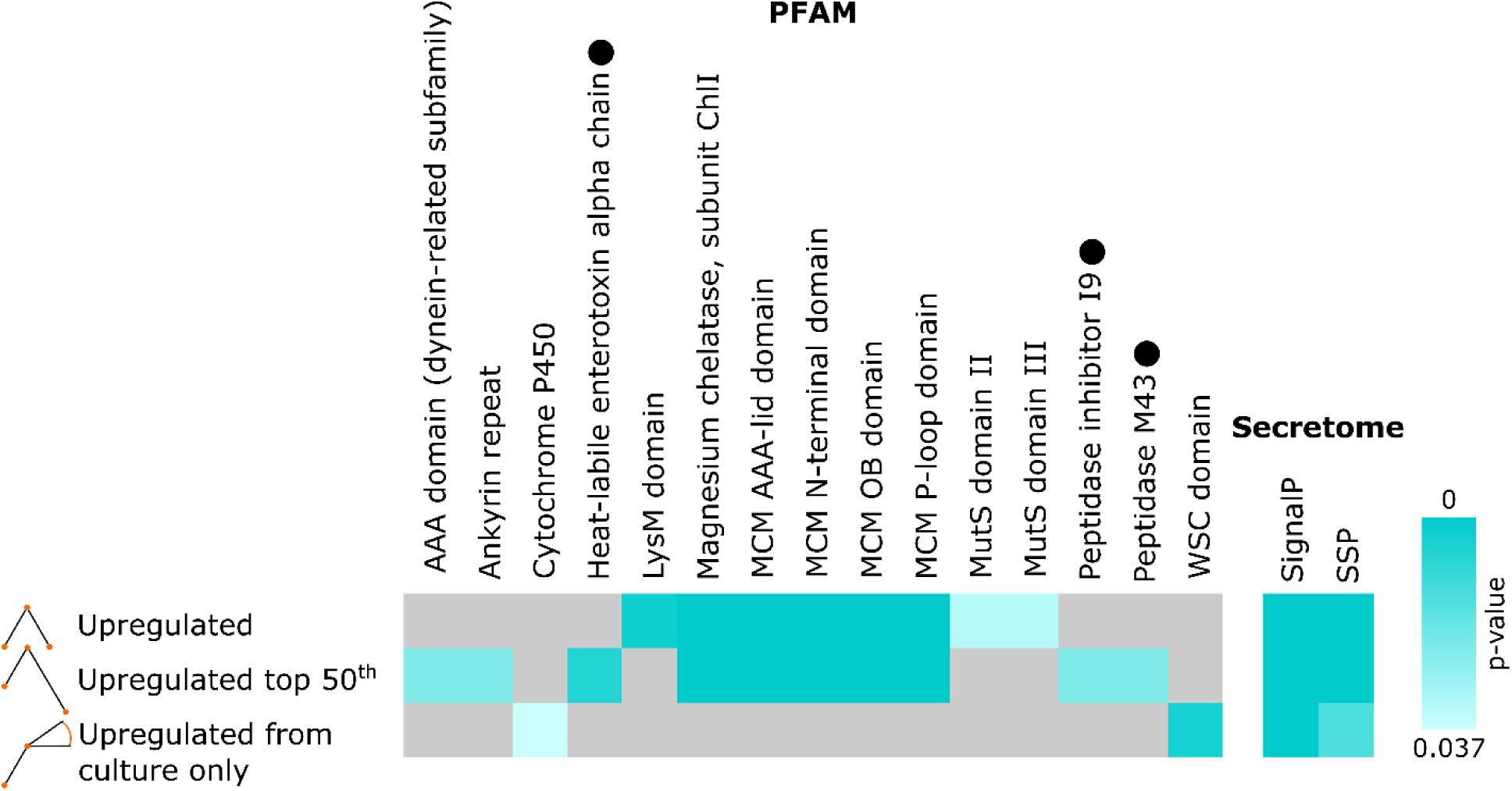
PFAM and secretion signal domain enrichment in DEG sets with increased transcription during live manipulation. Upregulated refers to genes with peak transcription during live manipulation. Upregulated top 50^th^ refers to the subset of upregulated genes with the strongest subsequent downregulation in dead manipulated samples. Upregulated from culture only are genes with increased transcription from culture to live manipulation, but then no change or increasing transcription with host death. PFAMs associated with genes that have possible roles in manipulation include: Heat-labile enterotoxin alpha chain, LysM domain, Peptidase inhibitor I9, and Peptidase M43 (indicated by black dots).

Notably, ant-infecting *Ophiocordyceps* genomes are enriched for heat-labile enterotoxins compared to generalist fungal pathogens and have been suggested to play a major role in *Ophiocordyceps* pathogenesis. For example, *O. kimflemingiae* has 36 putative enterotoxins, *Ophiocordyceps australis* has 20, and *Ophiocordyceps polyrhachis-furcata* has 22 (Wichadakul & al. 2015, de Bekker, Ohm, & al. 2017), while the generalist entomopathogens *C. bassiana, M. robertsii*, and *Isaria javanica* have 13, six, and five enterotoxins, respectively (Xiao & al. 2012, Lin & al. 2019, Mannino & al. 2019). The most strongly upregulated enterotoxin in *O. camponoti-floridani* displayed a >12,000-fold increase in transcripts from culture. Similarly, the *O. kimflemingiae* homolog to this gene displayed a marked > 3,000-fold upregulation (de Bekker & al. 2015). Additionally this putative enterotoxin is phylogenetically positioned in a clade containing only ant-manipulating *Ophiocordyceps* (de Bekker, OHM, & al. 2017) suggesting a specialized role in facilitating manipulation of ant hosts.

Other putative toxins were also upregulated in *O. camponoti-floridani* at the time of manipulation relative to growth in culture and in dead hosts. This included two Bordetella pertussis toxin A genes, which were also identified in the manipulation correlated fungal WGCNA module F1. Pertussis toxin enzymatic subunits of *Bordetella pertussis* carry amino acid sequence similarity in their active sites with heat-labile enterotoxins. As such, these ADP-ribosylating toxins also attack G protein-coupled receptor systems causing increased levels of cyclic AMP to accumulate (Locht & al. 1986). One of the putative pertussis toxins was also upregulated in *O. kimflemingiae* over the course of infection (i.e., 4-fold increase from culture to live manipulation in *O. camponoti-floridani*, and a 538-fold in *O. kimflemingiae*). The pertussis toxins in *O. camponoti-floridani* each carried an Ankyrin repeat domain, which contributed to the enrichment for this PFAM annotation (Fig.5).

Additionally, both *Ophiocordyceps* fungi upregulated a homologous secreted ribonuclease during manipulation that may function as a ribotoxin (i.e., 11-fold increase from culture to live manipulation in *O. camponoti-floridani*, 19-fold in *O. kimflemingiae*). We also found this ribotoxin in fungal manipulation WGCNA module F2. Unlike ADP-ribosylating toxins, fungal ribotoxins are considered ribonucleases with possible insecticidal effects (Olombrada & al. 2013).

#### Fungal Serine proteases are upregulated during manipulated biting behavior

Fungal subtilases are subtilisin-like serine proteases that have been implicated in entomopathogenic interactions. During infection, *Metarhizium anisopliae* hyphal bodies in insects produce increased levels of serine protease Pr1, leading to host death. Additionally, Pr1 overproducing strains of *M. anisopliae* decrease host-feeding and injections of Pr1 are toxic (St Leger & al. 1996). Often, these proteases contain a peptidase inhibitor I9 domain, which is present in propeptides and assists folding and activation once cleaved (reviewed in Figueiredo & al. 2018). The presence of an evolutionarily recent group of subtilases with unclear function that lack I9 domains may be associated with niche adaptation in the Entomophtoromycotina; fungi distantly related from *Ophiocordyceps*. Among the Entomophthoromycotina, the insect manipulating *Entomophthora muscae* and *Pandora formicae* indeed have I9-lacking subtilases, albeit not exclusively (Arnesen & al. 2018). A comparison of this protease group to subtilases of Ascomycetes demonstrated a notable dissimilarity. However, the representative of *Ophiocordyceps* in that analysis was a lepidopteran parasite outside the *O. unilateralis* species complex, *Ophiocordyceps sinesis* (Arnesen & al. 2018).

Inhibitor I9 domain encoding genes were found to be enriched among genes upregulated during manipulation (Fig.5). Two of these three I9 containing genes were subtilases with MEROPS S8A annotations similar to Pr1 found in *C. bassiana* and *M. anisopliae* (Joshi & al. 1995). In total, we found six S8A annotated *O. camponoti-floridani* genes upregulated during manipulation relative to culture. One of which was found in manipulation WGCNA module F1. Similarly, six serine proteases were upregulated during manipulation in *O. kimflemingiae* (de Bekker & al. 2015). Of the differentially expressed S8A subtilases in *Ophiocordyceps*, two lacked inhibitor I9 domains (one with 38-fold increase, the other increasing from 0 FPKM to 4 FPKM).

Expansion of protease gene copies has been suggested to be associated with a broader host range in fungal entomopathogens as a large repertoire of proteolytic enzymes may facilitate infection of various host species (Xiao & al. 2012, Lin & al. 2019). The ant manipulating *Ophiocordyceps* appear to be highly species-specific parasites (de Bekker, Quevillon, & al. 2014, Araújo & al. 2018, Sakolrak & al. 2018), and their relative number of subtilases fall in line with this suggested trend. The ant manipulating *Ophiocordyceps* carry fewer of these proteases (*O. camponoti-floridani* n = 16, *O. kimflemingiae* n = 18, and other *Ophiocordyceps* species n < 20) compared to the generalists (*C. bassiana* n = 43, *Cordyceps militaris* n = 26, *M. anisopliae* n = 55, and *Metarhizium acridum* n = 43) (Gao & al. 2011, Zheng & al. 2011, Xiao & al. 2012, de Bekker & al. 2015, Wichadakul & al. 2015). Which of these subtilases *Ophiocordyceps* species specifically employ for virulence in general or manipulation specifically is not clear without further study.

#### Modulation of ant insulin-like peptides and growth factors by fungal metalloproteases

Insulin/IGF signaling (IIS) pathways have been implicated in behavior, division of labor, and establishment of the reproductive caste in ants and bees (Ament & al. 2008, Chandra & al. 2018). Juvenile hormone and vitellogenin (VG) appear to play an important role in IIS pathways, although typically the strongest effects are observed during development and early life (Lengyel & al. 2006, Leboeuf & al. 2018, Opachaloemphan & al. 2018). Nutritional status, energy demands, and caste related behavior (e.g. foraging and exploration/locomotor activity) appear to be interlinked in these eusocial insects. Therefore, one possible scenario for parasitic disruption of host behavior is that fungal effectors could be targeting IIS pathways to modify host behavior mediated via changes in JH or VG.

Indeed, we found putatively secreted metalloprotease encoding genes upregulated during live manipulation that could be involved in infection and affect host IIS pathways. These genes carried PFAM PF05572|Peptidase_M43 annotations and additional MEROPS protease annotations. Genes with MEROPS M43.002 putatively function similarly to *mep1*, which assists fungi to counteract mammalian immune systems (Hung & al. 2005, Shende & al. 2018). Putative *mep1* genes in *O. camponoti-floridani* were upregulated during live manipulation relative to both culture and dead manipulated samples (Fig.5). One homologous *mep1* metalloprotease was also significantly upregulated during manipulation in *O. kimflemingiae* (i.e., 71-fold increase from culture to live manipulation in *O. camponoti-floridani*, 3-fold in *O. kimflemingiae*) (de Bekker & al. 2015). Other M43 annotations present in the *O. camponoti-floridani* genome predicted the presence of ulilysins (MEROPS M43.007) and pappalysins (MEROPS M43.004 or M43.005). Ulilysins and pappalysins are known to interact with IGF binding proteins that regulate levels of free IGF (Tallant & al. 2007). Two M43 metalloproteases carrying both ulilysin and pappalysin MEROPS annotations were upregulated from culture to manipulation in *O. camponoti-floridani* but not in *O. kimflemingiae*.

#### Ecdysteroid related genes in host manipulation

Ecdysteroids interact with networks involving JH, VG, and insulin-like peptides. As such, ecdysteroids play an indirect role in ant development and the establishment of reproductive castes (Libbrecht & al. 2013) that have distinct behavioral profiles. Ecdysteroids are also implicated in viral manipulation of caterpillars that display a summit disease phenotype. Baculovirus secretes an enzyme, EGT, which inactivates an ecdysone molting hormone and alters larval feeding behavior and development (O’Reilly 1995). In certain species of caterpillar, *egt* is implicated in driving fatal summit disease, while in others only altering pre-molting climbing behavior (Hoover & al. 2011, Han & al. 2015, Ros & al. 2015). Moreover, the entomopathogenic fungus *Nomuraea rileyi* appears to attack host ecdysteroid pathways by secreting an enzyme that disrupts normal host larval development (Kiuchi & al. 2003, Kamimura & al. 2012).

Possibly, ecydsteroids also play a role in manipulation by *O. camponoti-floridani* since we detected a phosphotransferase gene with eckinase and SignalP domains that was significantly higher expressed during manipulation than in culture (2-fold increase). This gene was also present in WGCNA module F2. Eckinase activity helps mediate the balance of active free ecdysteroids and inactive storage forms (Sonobe & al. 2006). As such, *O. camponoti-floridani* could be utilizing a similar strategy as baculovirus or *Nomuraea* to modify its host; by using an ecdysteroid modulating enzyme.

#### Upregulated protein tyrosine phosphatase implicated in insect hyperactivity and ELA

Also implicated in baculovirus infection, PTP has a suggested role in ELA phenotypes of infected caterpillars (Kamita & al. 2005, Katsuma & al. 2012), but not summiting (VAN HOUTE, ROS, & al. 2014). Notably, evidence from KATSUMA & al. (2012) indicated that enzymatic function of PTP is not necessary for behavioral changes in the host, which suggests a sole structural role for viral PTP that is necessary for successful infection. In contrast, work with a different baculovirus strain and host combination by VAN HOUTE & al. (2012) demonstrated that PTP enzymatic activity was indeed necessary for the induction of ELA and is hypothesized to act via changes in TO (VAN HOUTE & al. 2013).

What function PTP may have for fungal manipulation of its host is uncertain at this time. However, *O. camponoti-floridani* has seven putative *ptp* genes, of which five were upregulated from culture during manipulation, and three are putatively secreted. Moreover, four *ptp* genes were found in manipulation WGCNA modules F1 (n = 1) and F2 (n = 3) and two of these upregulated *ptp* genes in *O. camponoti-floridani* (365-and 8-fold increase from culture to manipulation) have homologs in *O. kimflemingiae* that were upregulated in similar fashion (28-and 2-fold increase) (de Bekker & al. 2015).

### Fungal secondary metabolites involved in manipulation and infection

We identified 25 secondary metabolite clusters and backbone genes in the genome of *O. camponoti-floridani* and determined the bioactive compound classes they produce from the annotated PFAM domains of their backbone genes. The genomes of *O. kimflemingiae* and *O. polyrhachis-furcata* contain comparable numbers of annotated clusters, 25 and 24, respectively (de Bekker & al. 2015, Wichadakul & al. 2015). Notably, 23 of the *O. camponoti-floridani* metabolite clusters contain genes with homologs identified previously in *O. kimflemingiae* as members of secondary metabolite clusters (de Bekker, Ohm, & al. 2017). Their gene expression patterns and functional annotations offer insights into the possible fungal secondary metabolites involved in manipulation and infection as discussed in more detail below.

#### Cluster 18: Manipulation-related aflatrem-like indole-diterpene alkaloid production

Metabolite cluster 18 putatively produces an alkaloid that appears to be a mycotoxin. The entirety of this cluster demonstrated a striking upregulation during infection (Fig. 6a), as did the homologous cluster in *O. kimflemingiae* (de Bekker & al. 2015). Additionally, six of eight cluster 18 genes were found in fungal manipulation WGCNA module F2. These clusters are highly similar in gene composition and organization between the two *Ophiocordyceps* species (Fig. 6b), with both this study and DE BEKKER & al. (2015) identifying one possible product to be an ergot alkaloid based on a TRP-DMAT backbone gene of the cluster (> 12,000 fold increase from culture to live manipulation in *O. camponoti-floridani*, 5,900 fold in *O. kimflemingiae*). These aromatic prenyltransferases are known to drive the first step of ergot alkaloid synthesis (Ding & al. 2008). This class of secondary metabolites are notorious mycotoxins with effects on animal physiology and behavior. Poisoning by ergot alkaloids such as lysergic acid (a LSD precursor), clavine alkaloids, or ergotamine can induce tremors, confusion, body temperature dysregulation, hallucinations, and other symptoms (Schiff 2006). Ergot alkaloids can additionally drive entomopathogenic virulence in fungi (Clay & Cheplick 1989, Panaccione & Arnold 2017).

**Figure 6.**
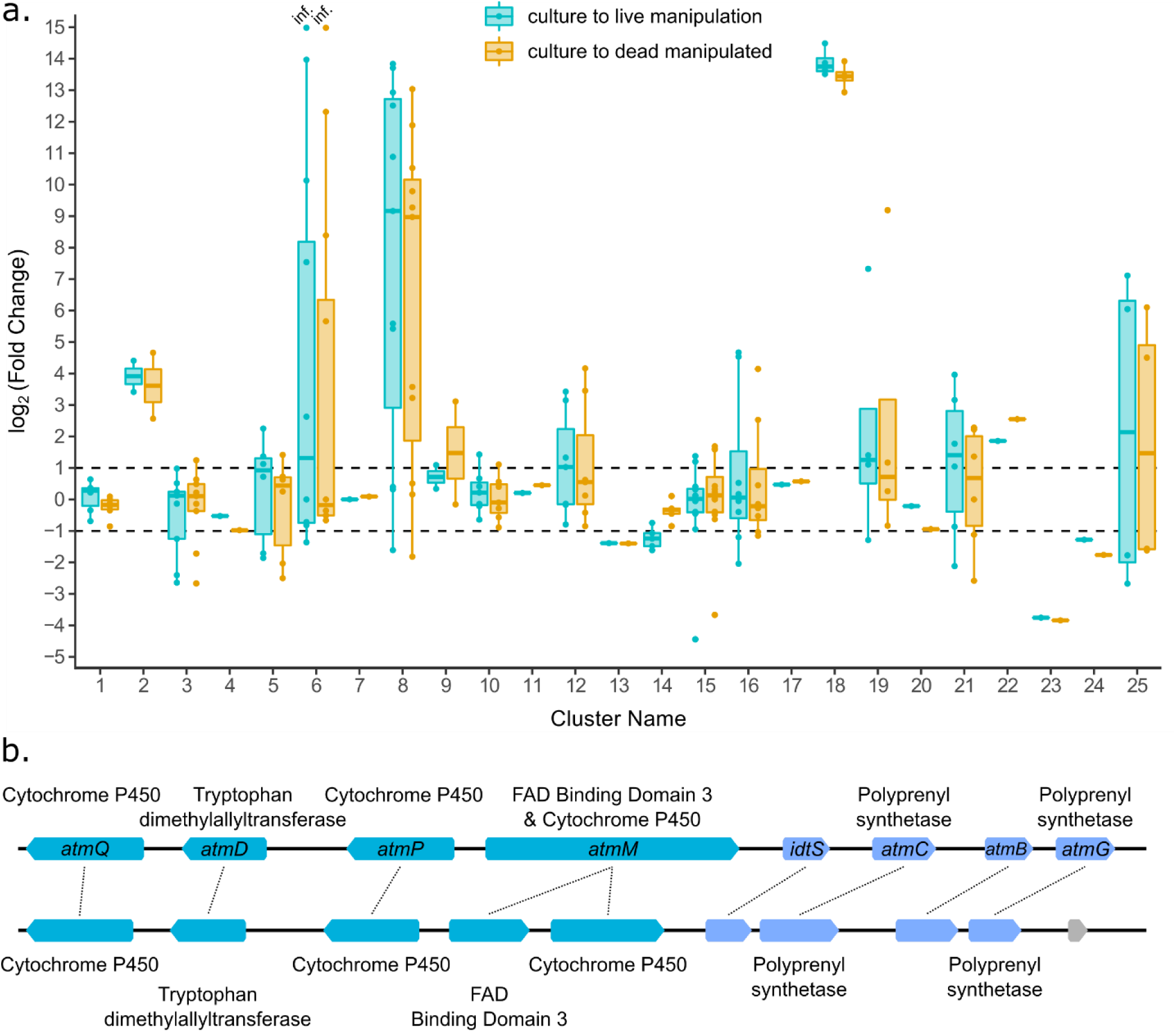
*(a.)* Fold change of secondary metabolite cluster genes transcripts during live-manipulation (blue) and in dead-manipulated samples (orange) relative to control culture. Points represent fold change of a single gene within a metabolite cluster. Dashed lines indicate a 2 fold change in transcript abundance (i.e. log_2_(Fold Change) = ±1). Clusters 2, 6, 8, 12, and 18 are possibly involved in pathways producing entomopathgenic compounds similar to aflatoxin, fusarin C, aflatoxin, citrinin, and aflatrem, respectively. Many genes of these clusters also displayed notable increases in transcript levels relative to culture. Points labeled “inf.” in cluster 6 have infinite fold increases due to culture FPKM values equal to 0. *(b.)* Schematic of cluster 18 genes (top) and corresponding homologs in *O. kimflemingiae* (bottom). PFAM annotations are labeled outside the depicted loci and, for *O. camponoti-floridani*, BLASTp hits for aflatrem synthesis proteins are indicated inside. Teal indicates cluster genes identified with our SMURF-based pipeline (Khaldi & al. 2010), blue are selected cluster-adjacent genes based on their similar expression, putative function, and proximity to the main cluster (not represented in *[a.]*). The locus in gray is unrelated to the cluster. Image modified from output generated by Geneious Prime (v 2019.0.3, Biomatters).

However, homology searches also indicated that cluster 18 may contribute to the production of an aflatrem-like indole-diterpene alkaloid (Fig. 6b). Aflatrem is a neurotoxic tremorgen that causes “stagger disease” in poisoned hosts and is closely related to other mycotoxins such as paxilline or lolitrem (Gallagher & Hawkes 1986). By interfering with big potassium (BK) channels, and, gamma amino butyric acid (GABA)-ergic and glutamatergic processes, aflatrem can induce muscle tremors, changes in activity level, and confusion (Valdes & al. 1985, Gant & al. 1987, Yao & al. 1989, Knaus & al. 1994). Furthermore, brain GABA levels have been shown to differ along caste lines in *C. floridanus* (Punzo & Williams 1994). The related indole-diterpene nodulisporic acid has also demonstrated insecticidal effects on multiple species (Smith & al. 2000, Shoop & al. 2001).

Adjacent to secondary metabolite genes identified by our SMURF-based pipeline (Khaldi & al. 2010) in cluster 18, we further identified putative cluster genes based on similar expression patterns, functional annotations related to secondary metabolism, and proximity (approximately within 3 kbp of another cluster gene, i.e., double the distance between any genes in the original cluster). Among these genes, we identified a similarly expressed terpene cyclase *atmB*-like gene adjacent to cluster 18. This gene has a homolog in *O. kimflemingiae* annotated as *paxB* (the corresponding paxilline synthesis gene of aflatrem), which suggests that aflatrem or a similar indole-diterpene is synthesized by cluster 18 rather than an ergot alkaloid (Fig. 6b).

We subsequently performed a BLASTp of translated *Aspergillus flavus* aflatrem synthesis genes (*atmA*, *atmB*, *atmC*, *atmD*, *atmG*, *atmM*, *atmP*, and *atmQ*) (Nicholson & al. 2009) against the entire *O. camponoti-floridan* genome. All proteins except AtmA scored BLASTp hits with the lowest E-value hits in or adjacent to cluster 18. All putative aflatrem synthesis homologs were present in a single cluster of the *O. camponoti-floridani* genome, while these genes are split between two genomic sites in *A. flavus* (Nicholson & al. 2009).

Despite not detecting any homologous proteins to AtmA in the *O. camponoti-floridani* genome, there is an upregulated gene in the cluster-adjacent set that has a BLAST annotation corresponding to *idtS*, but otherwise lacks a GO or PFAM annotation (Fig. 6b). Described in the lolitrem-producing fungi *Epichloë* and *Neotyphodium* as necessary for indole-diterpene synthesis (Schardl & al. 2013), the molecular role of IdtS is not entirely clear. Similarly, the role of AtmA in aflatrem synthesis has not been fully elucidated yet (Nicholson & al. 2009). The *O. camponoti-floridani idtS* homolog may also work in concert with the indole-diterpene synthesis genes identified in cluster 18, although we currently do not fully understand its function.

#### Clusters 8 and 2: Putative aflatoxin production during infection and manipulation

Aflatoxin is a potent mycotoxin and carcinogen with lethal effects on insects (Trienens & Rohlfs 2011). We predicted the polyketide metabolite clusters eight and two to produce an aflatoxin or similar metabolite, based on the PKS backbone gene in these clusters having a starter unit:acyl carrier protein transacylase (SAT) domain that facilitates the beginning step in the synthesis of aflatoxins (Brown & al. 1996). Upregulation of the majority of genes in putative aflatoxin-like clusters 8 and 2 during live manipulation suggests a role for this mycotoxin during the final stages of infection (Fig. 6a). Five genes from cluster 8 were also found in the manipulation associated WGCNA module F2. The PKS backbone of cluster 8 was found to be upregulated during manipulation in both *O. camponoti-floridani* and *O. kimflemingiae* (i.e., 43-fold increase from culture to live manipulation in *O. camponoti-floridani*, 4,350-fold in *O. kimflemingiae*). The PKS backbone of cluster 2 was also upregulated (11-fold increase from culture to live manipulation). However, cluster 2 homologs in *O. kimflemingiae* were downregulated during this time.

Using the *Aspergillus parasiticus* aflatoxin gene cluster (J. Yu & al. 2004), we performed a BLASTp search of 25 proteins against the *O. camponoti-floridani* genome. All but two genes (*aflI* and *aflX*) had at least one *O. camponoti-floridani* homolog. The PKS backbone of cluster 8 had homology to *aflC*, the PKS of the *A. parasiticus* cluster (E-value = 2.61e-98). Additionally, cluster 8 contained the top BLASTp hit for *aflJ*. Furthermore, *aflD* returned a low E-value hit within the cluster (1.94e-06). However, it fell short of our bit score 50 cutoff (46.21). Although these results do not demonstrate that cluster 8 produces aflatoxin, they do suggest it has a role in generating aflatoxin or similar compounds.

The PKS backbone of cluster 2 is also a putative homolog to AflC (E-value = 1.38e-93) and the O-methyltransferase was the top BLASTp hit for AflP, which generates the penultimate product in aflatoxin synthesis (J. Yu & al. 2004). Although our SMURF-based (Khaldi & al. 2010) approach only identified two genes in cluster 2 (a PKS and O-methyltransferase), similarly regulated adjacent genes could generate products consistent with the aflatoxin synthesis pathway. In proximity to cluster 2, the *O. camponoti-floridani* genome contains an upregulated gene encoding a protein homologous to AflQ, known to play an important role in the final step of aflatoxin synthesis (J. Yu & al. 2004). We also identified a possible homolog to *aflT* (BLASTp E-value = 4.22e-38), which is known to be involved in aflatoxin synthesis (J. Yu & al. 2004). A homolog of *A. parasiticus* Velvet-complex member *laeA* is also adjacent to the cluster, which functions as a regulator of the aflatoxin-intermediate sterigmatocystin (Bok & Keller 2004). The top BLASTp hit for AflR, which is a transcription factor involved in aflatoxin synthesis and interacts with LaeA in a regulatory feedback loop (Bok & Keller 2004, J. Yu & al. 2004) was found on the same contig, albeit distantly from cluster 2. Also possibly interacting with LaeA, two velvet domain containing genes were upregulated from culture to manipulation, one of which has a similarly upregulated homolog in *O. kimflemingiae*. The homologs of *alfC*, *alfP*, *alfT*, and a velvet transcription factor were also found in fungal WGCNA module F2.

Taken together, metabolite clusters 2 and 8 and their homologs in *O. kimflemingiae* suggest that both species of *Ophiocordyceps* have the capacity to produce an aflatoxin-like compound. However, contrasting transcriptomics data for cluster 2 across these species indicated that the production of aflatoxins during infection and manipulation might be differently regulated in these species.

#### Cluster 12: A citrinin cluster contributing to virulence

Cluster 12 is predicted to synthesize a compound similar to the polyketide citrinin. This mycotoxin is lethal to insects with nephrotoxic effects on Malpighian tubules, which perform kidney-like functions (Dowd 1989). Cluster 12 contains a backbone PKS that is a putative homolog to the citrinin PKS gene citS of *Monascus ruber* (BLASTp E-value = 4.49e-120) (He & Cox 2016). Additionally, genes in this cluster had high BLASTp matches to CitA and CitE that also participate in citrinin synthesis (E-value = 1.68e-60 and 1.56e-14 respectively). One cluster 12 gene was present in the manipulation correlated fungal WGCNA module F1 and two were in present in F2. However, no suitable matches were found within this cluster for the remaining cluster synthesis genes *citC*, *citD*, or *citB*. Although, these genes did have hits elsewhere in the genome. Of these putative citrinin synthesis homologs, only *citA* appeared differentially expressed, being upregulated during manipulation and host-death in both *O. camponoti-floridani* (Fig. 6a) and *O. kimflemingiae*. Possibly, cluster 12 genes are active and play an earlier role in infection than our sampling regime was able to capture.

#### Cluster 6: Fusarin C as a possible virulence or behavior modifying factor

Fusarin C is the likely product of cluster 6, consisting of nine genes, two of which were upregulated during live manipulation (Fig. 6a). Three other genes in this cluster were also present in WGNCA modules F1 (n = 1) and F2 (n = 2). Fusarin C is a carcinogenic mycotoxin and although produced by the entomopathogen *Metarhizium anisopliae*, insecticidal or antibacterial activity appears to be absent without culturing additives (Krasnoff & al. 2006). Additionally, insects possibly resist some effects of fusarin C by detoxification pathways that respond to xenobiotic and toxic challenges (Gelderblom & al. 1988, Gui & al. 2009, Rohlfs & Churchill 2011). However, fusarin C may have non-lethal effects as suggested by its apparent mycoestrogen activity, demonstrated in mammalian cells (Sondergaard & al. 2011). Estrogens could be involved in the production of ecdysteroids and VG with effects on development, reproduction, and diapause in insects (reviewed in Das 2016). Speculatively, exogenous estrogen activity from fusarin C produced by *Ophiocordyceps* could, thus, be disrupting normal host worker ant behavior and physiology by perturbing caste identity.

Four genes have been shown to be necessary for synthesis of fusarin C in *Fusarium fujikuroi*: *fus1*, *fus2*, *fus8*, and *fus9* (Niehaus & al. 2013). All four have homologs in cluster 6 and represented the top or only BLASTp result in the *O. camponoti-floridani* genome. Two additional fusarin C cluster proteins, Fus3 and Fus4 (Niehaus & al. 2013), had top BLASTp hits in cluster 6. Only *fus4* (seven-fold increase from culture to live manipulation) and *fus9* (0 FPKM to 2 FPKM) homologs were found to be upregulated during manipulation, while the other cluster genes were either not differentially expressed or downregulated. Homologs to all *F. fujikuroi* fusarin C genes were also detected in *O. kimflemingiae*, with four annotated in a secondary metabolite cluster. In this species, all but the *fus1* PKS-NRPS were upregulated during live manipulation relative to both culture and dead hosts (de Bekker & al. 2015).

Additionally, *Ophiocordyceps polyrhachis-furcata* contains the clustered homologs of the necessary fusarin C synthesis genes (Wichadakul & al. 2015). Similar to the case of aflatoxin, the presence of homologous fusarin C clusters among species of *Ophiocordyceps* suggests that all utilize fusarin C-like mycoestrogens, but when this metabolite is produced or used may differ between species. Possibly, fusarin C has a role earlier in infection when subtler behavioral changes such as altered locomotor activity manifest before manipulated clinging and biting. Changes in caste related behaviors, such as time spent foraging or nest occupation, may be plausible effects of the introduction of such a mycoestrogen and the underlying synthesis genes may no longer be strongly expressed by the final stage of manipulation in all *Ophiocordyceps*.

## Discussion

To shed light on possible mechanisms of host manipulation by ant infecting *Ophiocordyceps* fungi, we performed a *de novo* hybrid genome assembly of *O. camponoti-floridani* and RNAseq of both the fungal parasite and ant host *C. floridanus* before infection, during live manipulation, and after host death. We conducted our experiments and analyses in a comparative framework to highlight parallels to comparable manipulations observed in *O. kimflemingiae* – *C. castaneus* infections (de Bekker & al. 2015) that represent putative conserved or convergent mechanistic strategies used by the *Ophiocordyceps unilateralis* species complex (Fig. 7).

**Figure 7.**
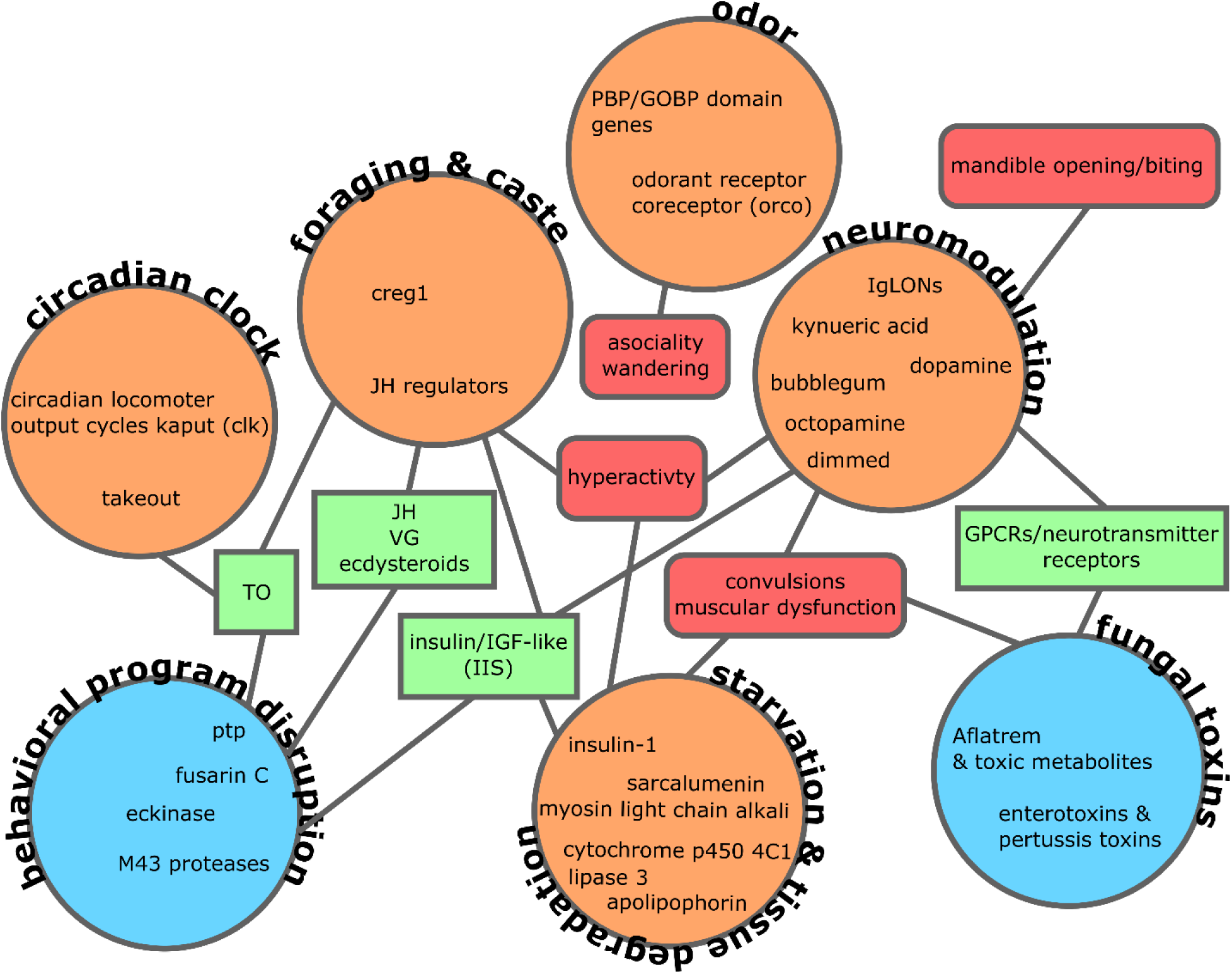
Major themes and candidate manipulation mechanisms emerging from comparative transcriptomics between *O. camponoti-floridani* – *C. floridanus* and *O. kimflemingiae* – *C. castaneus* interactions. Ant DEGS and proposed mechanisms (orange circles) are connected to fungal counterparts (blue circles) via shared molecular players (green rectangles) and phenotypes (red rounded-rectangles). These are scenarios we propose to be the most promising and intriguing with respect to manipulation, but do not exhaustively represent the available RNAseq data.

A growing body of literature has proposed possible fungal effectors of the ant-manipulating *Ophiocordyceps* based on genomic, transcriptomic, and metabolomic analyses (DE BEKKER, QUEVILLON, & al. 2014, de Bekker & al. 2015, Wichadakul & al. 2015, de Bekker, Ohm, & al. 2017, Kobmoo & al. 2018). Although phylogenetic and experimental evidence indicate that *Ophiocordyceps*-ant interactions are species specific (Kobmoo & al. 2012, de Bekker, Quevillon, & al. 2014, Araújo & al. 2018, Sakolrak & al. 2018), we hypothesized that *Ophiocordyceps* species share mechanisms to infect and manipulate their hosts as they face similar host and transmission related challenges (Chetouhi & al. 2015, Loreto & al. 2018). Indeed, we have found transcriptomic and genomic signals in *Ophiocordyceps camponoti-floridani* in line with previous work on other species, indicating several possible common contributors to infection and manipulation. Not only did these putative mechanisms of infection and manipulation emerge from comparisons across species (of both parasite and host), they were consistent among the analytical methods that we employed. By analyzing gene expression with PCAs, WGCNAs, and DEG identification, reoccurring genes and themes presented themselves. Additionally, numerous unannotated genes, including many SSPs, offer candidate manipulation genes and may serve as a wealth of hitherto novel effectors produced by the manipulating *Ophiocordyceps*.

In both the *O. camponoti-floridani* parasite and the *C. floridanus* ant host, we detected transcript abundances in line with previous work on *O. kimflemingiae* – *C. castaneus* interactions (de Bekker & al. 2015). However, some of our ant gene data and interpretation may be influenced by technical limitations. Firstly, we compared our ant RNAseq data to the ant dataset of DE BEKKER & al. (2015). In that work, the genome of *C. castaneus* was not available and transcripts were mapped to an earlier version of the *C. camponoti-floridani* genome than we use here (Bonasio & al. 2010). This inter-genome alignment may have reduced the resolution of the previously published *C. castaneus* RNAseq data. Secondly, the amount of RNA degradation and wide-spread dysregulation of gene expression in a recently deceased or moribund ant on the brink of death has likely introduced some noise into our ant data. These noise effects may have been exacerbated by the relatively low quantity of ant RNA reads remaining at the dead manipulated time point (ca. 4.9M reads). Despite these possible limitations, we still detected broad patterns, key genetic players, and well conserved responses to fungal infection and manipulation through our comparative transcriptomics approach (Fig. 7).

Corroborating observations that the ant manipulating *Ophiocordyceps* are highly specialized and often associate in a one fungus – one ant manner, we found relatively low number of putative subtilases in the genome of *O. camponoti-floridani* compared to generalist entomopathgenic fungi (Xiao & al. 2012, de Bekker, Quevillon, & al. 2014, Araújo & al. 2018, Sakolrak & al. 2018, Lin & al. 2019). Entomopathgenic fungi use subtilases to degrade insect cuticle (chitin) and, therefore, these enzymes can aid in host infection and killing. Secreted subtilases are upregulated in both *O. camponoti-floridani* and *O. kimflemingiae* during live manipulation of their ant hosts. Two of the six highly expressed subtilases in *O. camponoti-floridani* during manipulation lack I9 peptidase inhibitor domain annotations. Such subtilases have also been found in insect manipulating *Entomophthora* and *Pandora* and have been suggested to represent hallmarks of niche specialization (Arnesen & al. 2018). How subtilases would play a role in the mechanisms underlying host manipulation, however, remains to be investigated.

Also in line with previous works, laboratory infections of *C. floridanus* with *O. camponoti-floridani* induced manipulated biting and clinging behaviors synchronized by both time of day (i.e., ZT) and time after infection (i.e., dpi) (Hughes & al. 2011, de Bekker & al. 2015, Sakolrak & al. 2018). Unlike previous reports, however, manipulated ants in this study displayed clinging and biting before subjective dawn. In contrast, our preliminary field observations in central Florida indicated that manipulated biting in naturally infected *C. floridanus* can occur close to solar noon. This is similar to a previous field study in Thailand on *Ophiocordyceps camponoti-leonardi* (Hughes & al. 2011). These discrepancies between field and laboratory observations are probably due to altered environmental factors that function as zeitgebers, such as light, temperature, and humidity, in the laboratory compared to natural conditions.

A role for light-cues in the summiting aspect of *Ophiocordyceps* manipulation of ants has previously been proposed (Chung & al. 2017, Andriolli & al. 2018). Summit disease describes a behavioral manipulation phenomenon in which the host occupies locally elevated positions that are hypothesized to assist parasite transmission. Insect manipulating baculovirus strains also induce summiting behavior in silkworm hosts, and do so with an apparent phototactic element in coordinating manipulation (Kamita & al. 2005, VAN HOUTE, VAN OERS, & al. 2014, HAN & al. 2018). However, HAN & al. (2018) showed that light was required before but not during baculovirus induced summiting. Possibly, such a light coordinated behavior, but not direct phototaxis, underlies the pre-dawn summiting we observed in the laboratory. Although we have evidence for synchronization during manipulation across *Ophiocordyceps*-ant manipulations, the underlying cause and relatability between laboratory and natural conditions remain unclear.

The cooption and manipulation of circadian rhythms have been coined as the underlying mechanisms for synchronized biting and the disruption of caste specific foraging behaviors (DE BEKKER, MERROW, & al. 2014, de Bekker & al. 2015, de Bekker, Will, & al. 2017, de Bekker 2019). Ants display numerous behaviors under the control of their caste identities and circadian rhythms. Possibly, by disrupting these aspects of the host, related behaviors can be manipulated Fig. 7). Alterations in clock controlled behaviors may be affected by a dysregulation of the core clock protein CLK, as in both manipulated *C. camponoti-floridani* and *C. castaneus clk* was downregulated (de Bekker & al. 2015). A manipulated clock could also have led to the observed downregulation of a putative *to* homolog in both ant species. In turn, TO interacts with JH and affects foraging behavior (Sarov-Blat & al. 2000, Meunier & al. 2007, Schwinghammer & al. 2011, VAN HOUTE & al. 2013). Indeed, both JH activating and deactivating genes were differentially expressed during manipulation. As was CREG1, which is modulated by JH and has effects on cell growth that could be regulating caste development (Barchuk & al. 2007, Beckstead & al. 2007, Wurm & al. 2010). JH could, thus, be a mediator by which to change a variety of host behaviors since it interacts with many elements of ant physiology and development (Ament & al. 2008, Chandra & al. 2018, Opachaloemphan & al. 2018). Nevertheless, evidence in multiple species of ants indicate that JH has the most marked effects during critical periods of development (Opachaloemphan & al. 2018), not in adult ants. The possible effects of modifying JH dependent pathways in experienced adults (i.e., foragers exposed to *Ophiocordyceps* spores) in the context of fungal infection are unclear. However, as TO interacts with the circadian clock, JH, and foraging behavior, this could be a possibly enticing avenue for further exploration (SAROV-Blat & al. 2000, Meunier & al. 2007, VAN HOUTE & al. 2013).

One possible mechanism of parasitic disruption of TO-controlled behaviors that has been proposed in baculovirus is by way of PTP (VAN HOUTE & al. 2013). Upregulated *ptp* genes in *O. camponoti-floridani* and *O. kimflemingiae* are possibly implicated in driving aberrant host behavior as has been reported in baculovirus – caterpillar interactions (Kamita & al. 2005, VAN HOUTE & al. 2012, Katsuma & al. 2012, VAN HOUTE, ROS, & al. 2014). van Houte et al. (2013) also proposed PTP to act via interactions with the cGMP-dependent serine/threonine protein kinase Foraging (For), which underlies feeding behaviors in ants and other insects (Osborne & al. 1997, BEN-Shahar & al. 2002, BEN-Shahar & al. 2003, Lucas & Sokolowski 2009, Ingram & al. 2011) and serves as an intriguing candidate for fungal disruption. Circadian expression and phototatic effects of *for* have previously been shown (BEN-Shahar & al. 2003, Ingram & al. 2011). Therefore, if fungal PTP could dysregulate activity of For in ants, this would be a plausible strategy for the parasite to alter when and how long the ant host leaves the nest to engage in foraging behaviors and locomotor activity (Fig. 7).

Indeed, *Ophiocordyceps* infected ants have been reported to spend more time outside the nest compared to healthy nest mates (de Bekker, Quevillon, & al. 2014). Enhanced locomotor activity has been suggested to facilitate evacuating the nest and summiting (Hughes & al. 2011, de Bekker & al. 2015). Hyperactivity and ELA have also been reported in caterpillars infected by baculoviruses, and are a reoccurring theme in manipulative parasitisms (Kamita & al. 2005, Kathirithamby & al. 2010, VAN HOUTE, ROS, & al. 2014, de Bekker & al. 2018). This increased activity in infected ants may also be related to dysregulated foraging and feeding, as foraging workers necessarily are required to leave the nest and engage in exploratory behaviors. Such behaviors in ants are associated with caste, and perturbation of caste identity can generate atypical foraging patterns (Mersch & al. 2013, Simola & al. 2016). Caste-behavior, genetics, development, and nutritional status appear to be interlinked in eusocial insects (Ament & al. 2008, Libbrecht & al. 2013, Corona & al. 2016, Weitekamp & al. 2017, Chandra & al. 2018, Csata & Dussutour 2019), and fungal effectors that target pathways modulating caste related behaviors may underlie mechanisms of parasitic manipulation.

Pathways involving insulin-like peptides/IGF, estrogen-like compounds, and ecysteroids can modulate levels of JH and VG in insects, including carpenter ants and other eusocial hymenoptera, which could be possible targets for *Ophiocordyceps*. Altered JH and VG levels have been associated with developmental changes, changes in caste identity, and foraging behaviors (Bloch & al. 2000, Brent & al. 2006, Lengyel & al. 2006, Nelson & al. 2007, Ament & al. 2008, Velarde & al. 2009, Ament & al. 2010, Bonasio & al. 2010, Penick & al. 2011, Dolezal & al. 2012, Corona & al. 2013, Libbrecht & al. 2013, Corona & al. 2016, Das 2016, Chandra & al. 2018, Leboeuf & al. 2018). Evidence for fungal interference with these various aspects of host physiology came from multiple angles. Secreted M43 proteases produced by *O. camponoti-floridani* during manipulation could be interfering with IGF levels in the host (Tallant & al. 2007), and thereby play a role in disrupting host IIS pathways and related feeding behaviors. Paired with this effect, we also observed differential expression of a putative *insulin-1* gene in the host ant during manipulation (Fig. 7). If fusarin C acts as a mycoestrogen in *Camponotus* as it does with mammalian cells (Sondergaard & al. 2011), it could be dysregulating the production of ecdysterones and VG with possible effects on behavior (reviewed in Das 2016). Despite most reports indicating estrogen effects to be important during development, the upregulation of multiple members of a putative fusarin C metabolite cluster in *Ophiocordyceps* indicates that exogenous estrogen activity could disrupt caste identities and associated behavior of infected adult ants. Additionally, a putative secreted eckinase upregulated by *O. camponoti-floridani* during manipulation may modulate ant ecdysteroid levels and related feeding behaviors. Modifying titers of ecdysone molting hormones can alter larval feeding behavior and development, and is one of the proposed mechanisms underlying baculovirus manipulation of caterpillars (Hoover & al. 2011, Han & al. 2015, Ros & al. 2015).

In addition to attack on circadian clock elements, IIS and foraging pathways, and JH levels, ant neural tissues are plausibly dysregulated as part of behavioral manipulation by the fungal parasite (Fig. 7). Two ant gene modules, A14 and A15, were negatively correlated with live manipulation and characterized in part by large numbers of genes implicated in neuron development and neurotransmitter signaling. Furthermore, the dysregulation of putative *dimmed* or *bubblegum* homologs may result in changes in neuron integrity, secretory function, and downstream motor activity (Min & Benzer 1999, Hamanaka & al. 2010, Luo & al. 2013, Liu & al. 2016, Sivachenko & al. 2016). We also identified other differentially expressed genes that have putative neuromodulatory effects by affecting the synthesis of the neuroprotectant kynurenic acid, the production of dopamine, and responsiveness to octopamine. In each of these cases, *C. castaneus* differentially expressed homologs or similar genes during manipulation (de Bekker & al. 2015). A metabolomic study identified that the neuroprotectant ergothionine appears to be secreted by *O. kimflemingiae* during manipulation of *C. castaneus* (Loreto & Hughes 2019). Indeed, neural tissues appear to be among the last tissues to be severely degraded (Hughes & al. 2011, Fredericksen & al. 2017). Taken together, this suggests that the neural tissue is critical for fungal manipulation and may be partially preserved by compounds such as ergothionine or kynurenic acid while also being modified by fungal effectors operating via changes in ant biogenic amines and the disruption of neuron functions.

Many cellular receptors, including neurotransmitter receptors, are GPCRs that could serve as targets for fungal effectors. ADP-ribosylating toxins, such as heat-labile enterotoxin and pertussis toxin, cause detrimental accumulation of cyclic AMP in targeted cells by dysregulating GPCR signaling (Lin & al. 2010, Mangmool & Kurose 2011). Related toxins also suggest possible functions that *Ophiocordyceps* may employ such toxins for. Heat-stable enterotoxins appear to dysregulate pheromone production in insect fat bodies (Wiygul & Sikorowski 1986, Wiygul & Sikorowski 1991). Cytotoxic necrotizing factor-1, interacts with GTPases and contribute to *E. coli* invasion of central nervous tissues and crossing of the blood-brain-barrier in mammals (Khan & al. 2002). The ADP-ribosylating mosquitocidial toxin of *Bacillus spahericus* is similar to other bacterial toxins that act on G proteins and has lethal effects on mosquitoes (Thanabalu & al. 1991). Identified by both genomic and transcriptomic evidence, the secretion of enterotoxins and pertussis toxins could be a critical tool for *Ophiocordyceps* to invade, manipulate, and kill their ant hosts. Genomic evidence from the present study and previous ones revealed that enterotoxins are enriched in *Ophiocordyceps* genomes relative to generalist entomopathogens. Moreover, some enterotoxin gene clades group within the ant-manipulating species of *Ophiocordyceps* (de Bekker, Ohm, & al. 2017, Kobmoo & al. 2018). Additionally, multiple homologous enterotoxin and pertussis toxin genes were markedly upregulated in *O. camponoti-floridani* and *O. kimflemingiae* during manipulation (de Bekker & al. 2015). However, the exact function of heat-labile enterotoxins during pathogenesis remains elusive in entomopathogenic fungi (Mannino & al. 2019). Therefore, if these toxins indeed allow *Ophiocordyceps* fungi to interfere with host neurobiology to cause aberrant behaviors, or, invade certain tissues and promote cytotoxic effects, has yet to be determined.

An aflatrem-like mycotoxin produced as a secondary metabolite by *Ophiocordyceps* possibly underlies disease and manipulation symptoms such as convulsions (Hughes & al. 2011), muscle hypercontraction (Mangold & al. 2019), and atypical exploratory behaviors (de Bekker, Quevillon, & al. 2014). By disrupting neuron activity, aflatrem can induce muscle tremors, changes in activity level, and discoordination in mammals (Valdes & al. 1985, Gallagher & Hawkes 1986, Gant & al. 1987, Yao & al. 1989, Knaus & al. 1994). Aflatrem gene cluster synteny and upregulation during manipulation are well conserved between *O. camponoti-floridani* and *O. kimflemingiae*, suggesting an important role for this mycotoxin in regulating manipulation. Metabolite clusters are suggested to reflect ecological and evolutionary pressures and allow insight into possible critical functions for the life history of a fungus (Slot & al. 2019). Possible fitness benefits are suggested by these genes being clustered at a single genomic site and their high degree of conservation between *O. camponoti-floridani* and *O. kimflemingiae*. Such organization may allow these fungi to better regulate aflatrem genes in tandem and maintain them as a physically linked genetic unit, unlike *A. flavus* for example, which carries these genes at two distinct sites.

Possibly related to the effects of fungal toxins, manipulated *Camponotus* also suffered a dysregulation of dopamine and octopamine signaling, which have neuromodulatory effects in invertebrates. Disruption of the function of these amines has been linked to foraging and hyperactivity in insects (David & Verron 1982, Schulz & al. 2002, reviewed in Yamamoto & Seto 2014, Yang & al. 2015), and changing activity of such neuromodulators in the ant could feasibly contribute to observed ELA and hyperactivity preceding manipulated biting. Moreover, venom-induced hypokinesia in cockroaches has been linked to modulation of octopamine levels, most likely through manipulation of octopamine receptors (Libersat & Gal 2014). Dopamine has also been found to induce opening of mandibles and biting behavior in ants, a function indispensable for the “death grip” observed in manipulated ants (Szczuka & al. 2013). Although both *C. floridanus* and *C. castaneus* share an upregulation of a putative dopamine synthesis gene, octopamine reception is down in the former and up in the latter. This indicates that, if octopamine is critical to host manipulation, the specific octopamine-involved mechanisms may differ across the ant-manipulating *Ophiocordyceps* (de Bekker & al. 2015). Octopamine can also modulate insulin levels in insects (Li & al. 2016), which could drive behavioral responses normally linked to nutritional state. We predict that *C. floridanus* and *C. castaneus* ants are starving by the time the fungus has taken control of their behavior. Indeed, we found differentially expressed *C. floridanus* and *C. castaneus* genes related to nutrition and metabolism that corroborate this. In addition, the downregulation of odor receptors that we found across both host ant species might help ensure starvation-induced hyperactivity is not suppressed during the late stages of manipulation.

Odor detection and discrimination is crucial for proper social organization in eusocial insects, including ants. The large number of odorant receptors and OBPs in ant species as compared to bees and other Hymenopterans is an indicator of the complex chemical communication that underlies ant societies. The differential expression of genes encoding OBPs, especially PBPs, appeared to be a consistent theme in both *C. floridanus* and *C. castaneus* infected by *Ophiocordyceps*, suggesting that dysregulation of odorant receptors and OBPs contribute to the desensitization of olfaction and wandering behaviors that precede manipulated biting (de Bekker & al. 2015). The highly conserved odorant receptor, *orco*, regulates sensory neuron activity and odorant responses (Stengl & Funk 2013). Knockdown of *orco* expression has resulted in a significant decrease in antennal electrophysiological response and a reduction of sensitivity to semiochemicals in moths (Lin & al. 2015) and true bugs (Zhou & al. 2014), respectively. Ant *orco* mutants show a general reduction in odor sensitivity and reduced ability to forage or detect prey (Yan & al. 2017). Resulting in an environmentally and socially less receptive phenotype, dysregulation of *orco* expression may be involved the characteristic wandering behavior of *Ophiocordyceps* manipulated ants that precedes the death grip (Hughes & al. 2011, de Bekker & al. 2015). Indeed, *orco* was differentially expressed in both *C. floridanus* and *C. castaneus* which suggests that *Ophiocordyceps* affects the sensitivity of odor detection in its manipulated host. However, *orco* was up-regulated in *C. floridanus*, while its homolog was down-regulated in *C. castaneus*. As with potential octopamine-regulated mechanisms, manipulating strategies involving ORCO might, thus, differ across *Ophiocordyceps* species.

Taken together, we propose a number of fungal candidate manipulation genes and possible behavioral pathway responses in the ant that could be driving the manipulated climbing, biting, and clinging behaviors observed in *Ophiocordyceps*-manipulated individuals. The fact that we find many candidate genes across two independently performed infection experiments with two different host and parasite species provides strong evidence for the involvement of these genes. However, the candidates that we identified by comparing two species-specific manipulation events must still be functionally tested for a more precise understanding of how, and if, they play a critical role in the manipulation of host behavior. Similarly, follow-up metabolomic approaches would validate the production of secondary metabolites by the putative gene clusters we propose here. An analysis of possible protein-level mimicry by the parasite to interfere with host processes and physiology may be fruitful as well (Hebert & al. 2015). Of particular interest may be the comparison of multiple *Ophiocordyceps* and their respective ant hosts to identify possible species-specific mimicry that could indicate tight coevolutionary relationships and precise mechanisms of manipulation. Such studies could eventually be expanded to other parasitic manipulation systems since we find signatures of potentially convergently evolved mechanisms across manipulators. As such, the findings of this study provide an important springboard towards deeper functional and evolutionary understandings of the molecular mechanisms underlying host manipulation. In turn this might also offer insights into novel bioactive compounds and the neurobiology of animal behavior in general.

**Table 1.** O. camponoti-floridani genome assembly.

| Assembly characteristic | Value | Annotation | Number of genes |
| --- | --- | --- | --- |
| contigs | 13 | BLAST | 6291 |
| size (Mbp) | 30.5 | PFAM | 5460 |
| N50 (Mbp) | 3.8 | GO | 3515 |
| largest contig (Mbp) | 5.6 | SignalP | 801 |
| predicted genes | 7455 | SSP | 271 |
| BUSCO% | 99.7 | TMHMM | 1372 |
| GC% | 48.4 | 2° metabolism | 111 |
|  |  | MEROPS | 243 |
|  |  | transcription factor | 206 |

## Supporting information

Ant_RNAseq

Fungal_RNAseq

## Supplementary data

Available on bioRxiv.

## Acknowledgments

We thank the Laboratory for Functional Genome Analysis at the Ludwig Maximilians Universität for sequencing support. We also thank Hannah Christensen for isolating the Arb2 fungus, and Andrea Bender, Sara Linehan, and Brianna Santamaria for assistance with ant observations.

## Author contributions

IW, BD, TT, CdB wrote the manuscript, ran the infection experiments, and analyzed data; IW and AB generated sequence data; IW and RA assembled and annotated the genome; CdB and IW conceived the study.

## Data availability

Genome assembly and RNAseq data sets will be uploaded to NCBI

