## Supplemental Figures & Tables for "Genetic underpinnings of host manipulation by *Ophiocordyceps* as revealed by comparative transcriptomics"

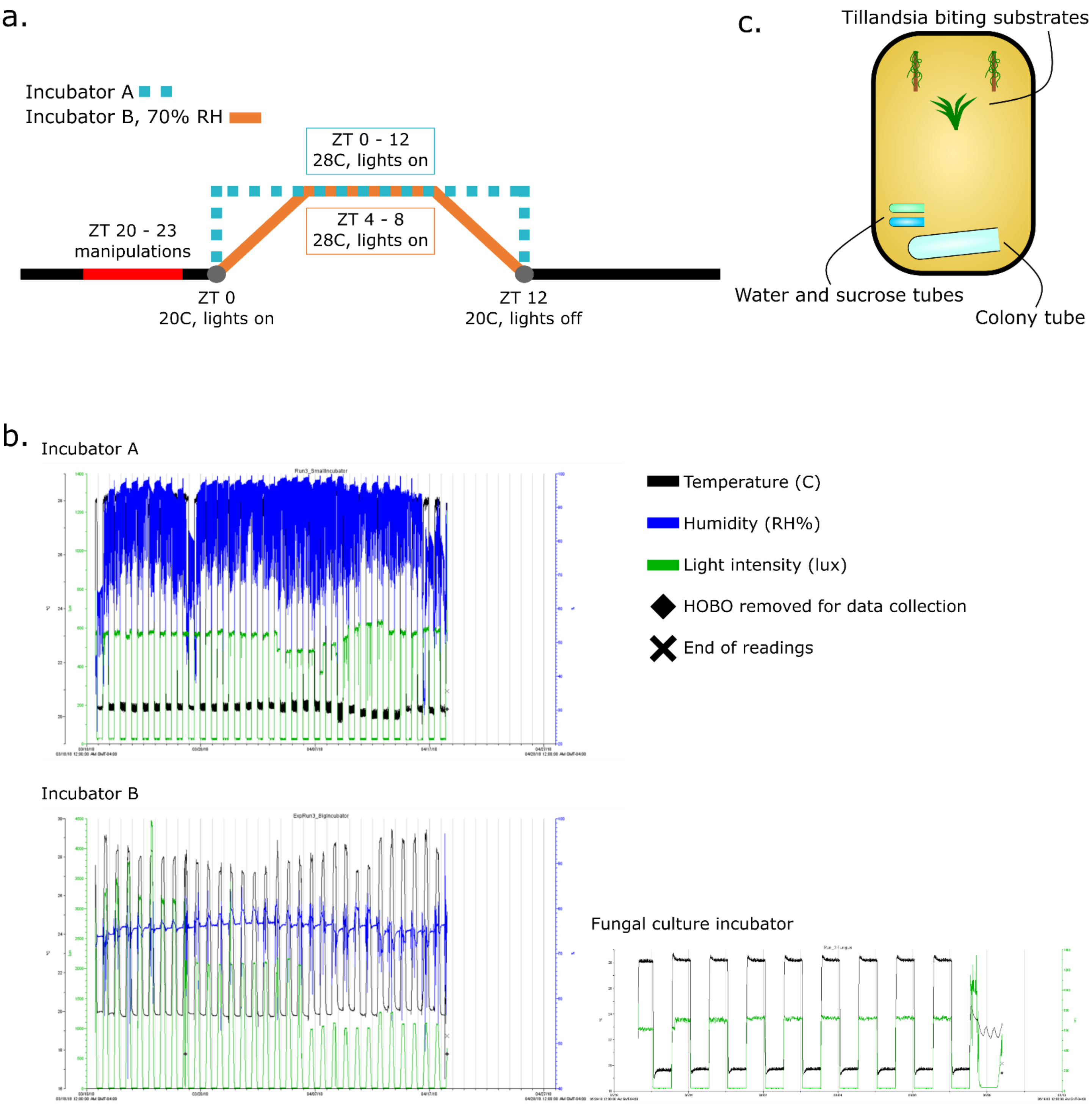


*Supplementary figure 1. (a.)* Incubator climate cycles. Incubator B included a constant humidity at 70% RH. *(b.)* HOBO data for duration of the experiment of incubator A, B, and the one used to culture fungi. *(c.)* Schematic of ant infection experiment enclosure, 33 cm x 22 cm.

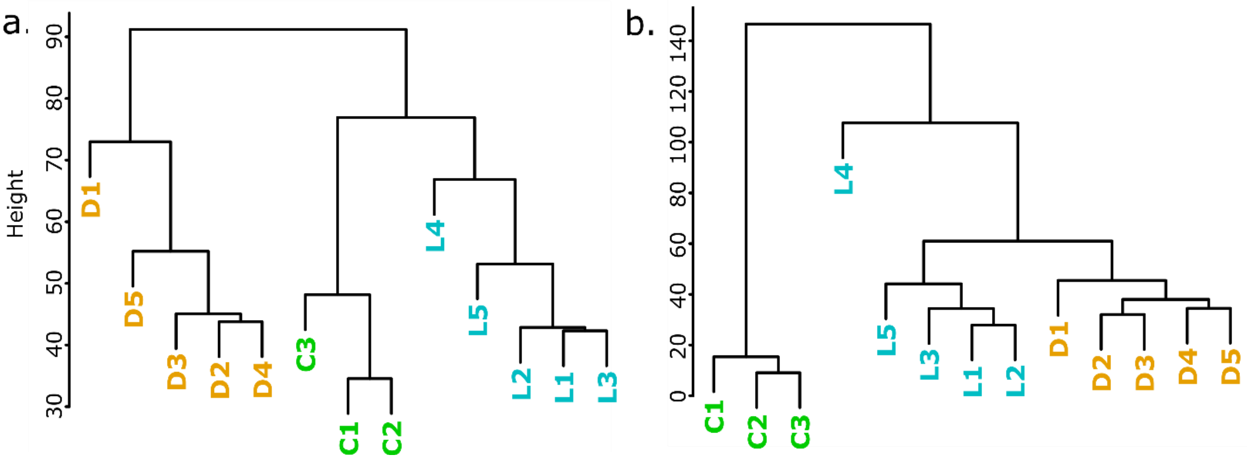


*Supplementary figure 2.* Unsupervised dendrogram clustering of biological replicates based on normalized gene expression levels (FPKM). Control (green, C1 – C3), live manipulation (blue, L1 – L5), and dead manipulated (orange, D1 – D5) samples largely cluster into their defined biological groups. *(a.)* Dendrogram of *C. floridanus* transcriptome data. The clustering of replicates indicates that living ants, whether healthy or manipulated, are more similar to each other than to recently expired ants after manipulation. *(b.)* Dendrogram of *O. camponoti-floridani* transcriptome data. The clustering of replicates indicates that fungi that actively interact with their host have more similar gene expression profiles to each other than to fungal growth under pre-infection conditions.

*Supplementary table 1.* Ant PC1 top 20 genes, ranked by highest to lowest loading value (0.099 – 0.063). For every gene, normalized expression values were highest in control samples and lowest in dead manipulated samples. Column "Homolog" refers to presence of a homolog between the current C. floridanus assembly and proteins annotated in DE BEKKER & al. (2015) with corresponding RNAseq data displaying similar patterns our study and DE BEKKER & al. (2015). Comments related to putative functions from Uniprot (BATEMAN 2019) and Interpro (FINN & al. 2017).

| BLAST | PFAM | Homolog | Comments |
|---|---|---|---|
| uncharacterized LOC105257597 | PF00026\|Eukaryotic aspartyl protease, PF00026\|Eukaryotic aspartyl protease | no | Lysosome activity |
| protein CREG1 | PF13883\|Pyridoxamine 5'-phosphate oxidase | no | Control of cell growth and apoptosis |
| lysosomal aspartic protease | PF00026\|Eukaryotic aspartyl protease | yes | Lysosome activity |
| myogenesis-regulating glycosidase-like | PF01055\|Glycosyl hydrolases family 31 | no | Muscle related via interaction with IGF2, jaw adductors known to be hypercontracted and degraded |

| | | | |
|---|---|---|---|
| lysosomal aspartic protease | PF00026\|Eukaryotic aspartyl protease | no | Lysosome activity |
| NA | NA | no | |
| alpha-amylase 1 | PF00128\|Alpha amylase, catalytic domain, PF02806\|Alpha amylase, C-terminal all-beta domain | no | Starch catabolism, may suggest reduction in food intake |
| venom acid phosphatase Acph-1 | PF00328\|Histidine phosphatase superfamily (branch 2) | no | Possibly a non-venom acid phosphatase associated with lysosomes, as *acph-1* is associated with venom glands absent in ant heads |
| NA | NA | no | |
| regucalcin | PF08450\|SMP-30/Gluconolactonase/LRE-like region | no | Modulates calcium dependent processes and signaling |
| alpha-amylase A-like | PF02806\|Alpha amylase, C-terminal all-beta domain, PF00128\|Alpha amylase, catalytic domain | yes | Starch catabolism, may suggest reduction in food intake |
| uncharacterized LOC105253145 | PF00135\|Carboxylesterase family | no | |
| venom acid phosphatase Acph-1-like | PF00328\|Histidine phosphatase superfamily (branch 2) | no | Possibly a non-venom acid phosphatase associated with lysosomes, as *acph-1* is associated with venom glands absent in ant heads |
| uncharacterized LOC105250203 | PF07464\|Apolipophorin-III precursor (apoLp-III) | yes | Fat body and lipid regulation, possibly link to starvation/hunger signaling |
| NA | NA | no | |
| pheromone-binding protein Gp-9-like | PF01395\|PBP/GOBP family | no | Odorant binding |
| troponin C-like | PF13499\|EF-hand domain pair; PF13833\|EF-hand domain pair | no | |
| cytochrome P450 4g15-like | PF00067\|Cytochrome P450 | no | Steroid synthesis |

| | | | |
|---|---|---|---|
| actin, muscle | PF00022\|Actin | no | Muscle related, jaw adductors known to be hypercontracted and degraded |
| uncharacterized LOC105258464 | PF01395\|PBP/GOBP family | no | Odorant binding |

*Supplementary table 2.* Ant PC2 top 20. 0.098 - 0.054. Column “Homolog” refers to presence of a homolog between the current C. floridanus assembly and proteins annotated in DE BEKKER & al. (2015) with corresponding RNAseq data displaying similar patterns our study and DE BEKKER & al. (2015). Comments related to putative functions from Uniprot (BATEMAN 2019) and Interpro (FINN & al. 2017).

| BLAST | PFAM | Homolog | Comments |
|---|---|---|---|
| NA | NA | no | |
| NA | NA | no | |
| uncharacterized LOC105253145 | PF00135\|Carboxylesterase family | no | |
| uncharacterized LOC105254189 | PF13639\|Ring finger domain | yes | |
| cytochrome P450 9e2 | PF00067\|Cytochrome P450 | no | |
| sialin | NAPF07690\|Major Facilitator Superfamily | yes | |
| uncharacterized LOC105251735 | NA | yes | |
| pheromone-binding protein Gp-9-like | PF01395\|PBP/GOBP family | no | Odorant binding |
| NA | NA | no | |
| uncharacterized LOC112639471 | PF15430\|Single domain von Willebrand factor type C | no | Response to infection and changes in nutritional status, especially in arthropods. |
| protein CREG1 | PF13883\|Pyridoxamine 5'-phosphate oxidase | no | Control of cell growth and apoptosis |
| defensin | PF01097\|Arthropod defensin | no | Arthropod immunity, especially against bacteria |
| uncharacterized LOC112639414 | PF01395\|PBP/GOBP family | no | Odorant binding |

| | | | |
|---|---|---|---|
| 2-amino-3-ketobutyrate coenzyme A ligase, mitochondrial-like | PF00155\|Aminotransferase class I and II | no | |
| uncharacterized LOC105255569 | PF13445\|RING-type zinc-finger, PF00240\|Ubiquitin family | no | |
| regucalcin | PF08450\|SMP-30/Gluconolactonase/LRE-like region | no | Modulates calcium dependent processes and signaling |
| ankyrin-1 | PF07525\|SOCS box, PF00023\|Ankyrin repeat, PF12796\|Ankyrin repeat | yes | |
| protein CREG1 | PF13883\|Pyridoxamine 5'-phosphate oxidase | yes | Control of cell growth and apoptosis |
| cytochrome P450 4C1-like | PF00067\|Cytochrome P450 | no | JH interacting, starvation response (LU & al. 1999) |
| WEB family protein At4g27595, chloroplastic | NA | no | Most BLAST results lacked meaningful annotations |

*Supplementary table 3.* Fungal PC1 top 20 genes, ranked by highest to lowest loading value (0.084 – 0.062). All have homologs in O. kimflemingiae with peak FPKM during live manipulation. Comments related to putative functions from Uniprot (BATEMAN 2019) and Interpro (FINN & al. 2017).

| BLAST | PFAM | Secretion | Comments |
|---|---|---|---|
| MFS transporter | PF07690\|MFS_1 | | Small solute transporter |
| protein kinase domain protein | | SignalP, SSP | |
| aromatic prenyl transferase | PF11991\|Trp_DMAT | | Cluster 18 (aflatrem) |
| GPI anchored serine-rich protein | | | |
| G-protein coupled receptor protein | PF00002\|7tm_2 | | |
| SCP-like extracellular protein | PF00188\|CAP | SignalP | Cysteine-rich secretory proteins, antigen 5, and pathogenesis-related 1 protein |

| | | | |
|---|---|---|---|
| alpha-ketoglutarate-dependent taurine dioxygenase | PF02668\|TauD | | |
| putative enterotoxin | PF01375\|Enterotoxin_a | SignalP | Highly expressed and most homologous enterotoxin |
| amidohydrolase family protein | PF04909\|Amidohydro_2 | | |
| P450 monooxygenase | PF00067\|p450 | | Cluster 18 (aflatrem) |
| prenyl transferase | PF00348\|polyprenyl_synt | | Cluster 18 (aflatrem) |
| P450 monooxygenase | PF01494\|FAD_binding_3, PF00067\|p450 | | Cluster 18 (aflatrem) |
| beta-lactamase family protein | PF11954\|DUF3471, PF00144\|Beta-lactamase | | Antibiotic resistance |
| P450 monooxygenase | PF00067\|p450 | | Cluster 18 (aflatrem) |
| hypothetical protein CDD80_6012 | | SignalP, SSP | |
| hypothetical protein CDD80_6620 | | SignalP, SSP | |
| tyrosinase 2 | PF00264\|Tyrosinase | SignalP | Associated with melanin production, stress and immune interactions |
| carbohydrate-binding module family 19 protein | | | |
| AtmB protein | | | Cluster 18 (aflatrem) |
| Pyruvate/Phosphoenolpyruvate kinase | PF13714\|PEP_mutase, PF00463\|ICL | | |

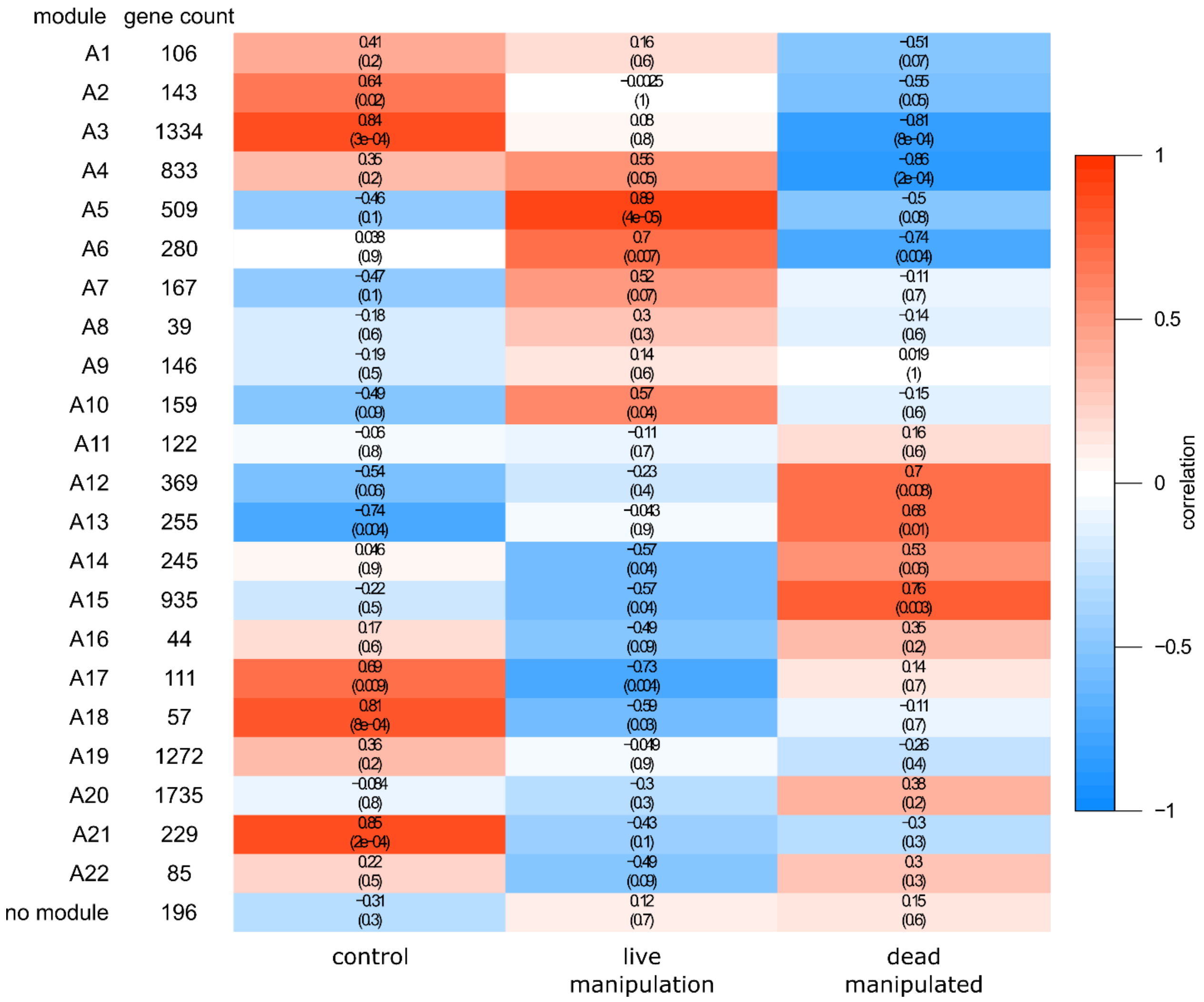


*Supplementary figure 3.* WGCNA of ant normalized gene expression data correlated to sample type. Correlation values are the top value in each cell, the p-values are in parentheses below.

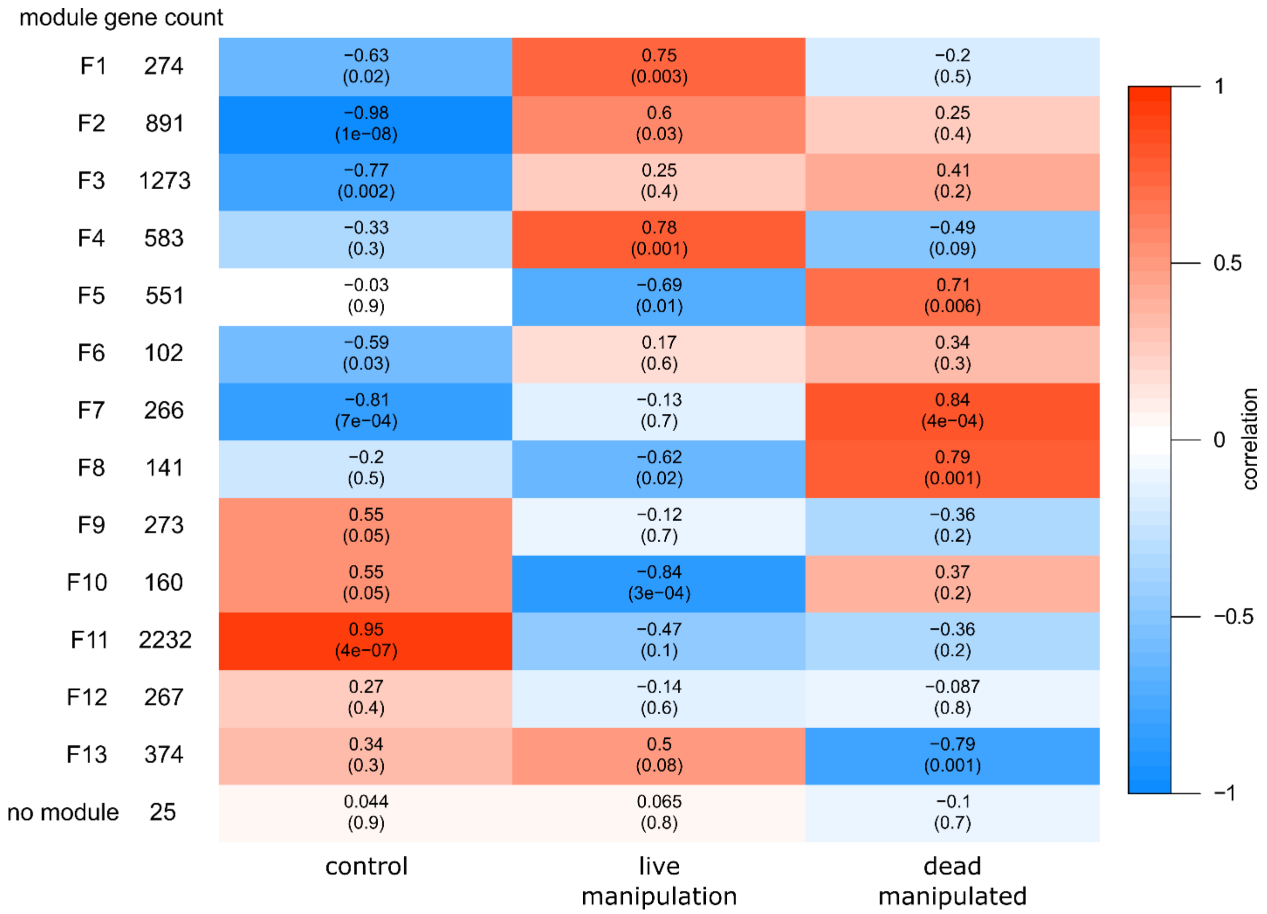


*Supplementary figure 4.* WGCNA of fungus normalized gene expression data correlated to sample type. Correlation values are the top value in each cell, the p-values are in parentheses below.

*Supplementary table 4.* Ant genes leading to enrichment of the rhodopsin family 7 transmembrane PFAM domain in ant WGCNA modules A14 and A15. Some of these genes are putatively involved in light sensing as rhodopsin, but many BLAST annotations seemingly indicate diverse cell signaling functions related to neurotransmitters. 5-hydroxtryptamine is synonymous with serotonin.

| BLAST annotation |
|---|
| 5-hydroxytryptamine receptor 1 |
| 5-hydroxytryptamine receptor 2A |
| 5-hydroxytryptamine receptor 2B |
| adenosine receptor A2b |
| allatostatin-A receptor |

| |
|---|
| alpha-2A adrenergic receptor |
| cholecystokinin receptor type A |
| dopamine D2-like receptor |
| dopamine receptor 1 |
| dopamine receptor 2 |
| G-protein coupled receptor 52 |
| lutropin-choriogonadotropic hormone receptor |
| melatonin receptor type 1B |
| muscarinic acetylcholine receptor DM1 |
| neuropeptide CCHamide-2 receptor |
| neuropeptide FF receptor 1 |
| neuropeptides capa receptor |
| octopamine receptor beta-1R |
| octopamine receptor beta-3R |
| opsin, ultraviolet-sensitive |
| probable G-protein coupled receptor B0563.6 |
| pyrokinin-1 receptor |
| somatostatin receptor type 2 |
| tachykinin-like peptides receptor 99D |
| trace amine-associated receptor 2-like |
| tyramine receptor 1 |
| uncharacterized |
| uncharacterized |
| uncharacterized |

*Supplementary table 5.* Ant genes leading to enrichment of immunoglobulin PFAM domains in ant WGCNA modules A14 and A15. Many of these genes have developmental and functional associations with neuronal tissues (but not always exclusively).

| BLAST annotation | Neuron associated | References |
|---|---|---|

| | | |
|---|---|---|
| basement membrane-specific heparan sulfate proteoglycan core protein | yes | (LINDNER & al. 2007, CHO & al. 2012) |
| cell adhesion molecule 2 | - | |
| Down syndrome cell adhesion molecule-like protein Dscam2 | yes | (CLANDININ & ZIPURSKY 2002, LAH & al. 2014) |
| follistatin-related protein 5 | yes | (BICKEL & al. 2008, PENTEK & al. 2009) |
| hemicentin-2 | - | |
| igLON family member 5 | yes | (CARRILLO & al. 2015) |
| immunoglobulin domain-containing protein oig-4 | yes | (RAPTI & al. 2011, SENGUPTA & al. 2019) |
| inactive tyrosine-protein kinase 7 | - | |
| irregular chiasm C-roughest protein | yes (esp. antennae) | (RAMOS & al. 1993, VENUGOPALA REDDY & al. 1999) |
| lachesin | yes | (KARLSTROM & al. 1993, STRIGINI & al. 2006) |
| leucine-rich repeat-containing protein 24 | yes | (DOLAN & al. 2007, CARRILLO & al. 2015) |
| leucine-rich repeat-containing protein 4 | yes | (DOLAN & al. 2007, CARRILLO & al. 2015) |
| neogenin | yes | (WILSON & KEY 2007) |
| nephrin | yes (mammal) nephrocytes (insect) | (ZHUANG & al. 2009, LI & al. 2011) |

| | | |
|---|---|---|
| netrin receptor UNC5C | yes | (KELEMAN & DICKSON 2001, KANG & al. 2019) |
| neuroglian | yes | (GODENSCHWEGE & MURPHEY 2009, GOOSSENS & al. 2011) |
| neurotrimin | yes (mammal) | (KRIZSAN-AGBAS & al. 2008, SANZ & al. 2015) |
| neurotrimin-like | yes | (KRIZSAN-AGBAS & al. 2008, SANZ & al. 2015) |
| obscurin | -<br>(muscle) | (KATZEMICH & al. 2012, KATZEMICH & al. 2015) |
| opioid-binding protein/cell adhesion molecule homolog | yes (mammal) | (MIYATA & al. 2003, REED & al. 2007) |
| peroxidasin | - | |
| protein borderless | yes | (SHAW & al. 2019) |
| protogenin | yes | (WONG & al. 2010, YU & al. 2013) |
| T-lymphocyte activation antigen CD86 | - | |
| tyrosine-protein phosphatase Lar | yes | (SETHI & al. 2010, AGRAWAL & HARDIN 2016) |
| zwei Ig domain protein zig-8 | yes | (BÉNARD & al. 2012, CHENG & al. 2019) |

*Supplementary table 6.* Putatively secreted proteins are transcribed during manipulation, but many genes lack functional PFAM domain annotation.

| | SignalP | | SSP | |
|---|---|---|---|---|
| | **Total** | **PFAM** | **Total** | **PFAM** |
| **Genome** | 801 | 409 | 271 | 52 |
| **Upregulated** | 77 | 39 | 31 | 6 |
| **Upregulated from culture** | 239 | 129 | 85 | 19 |
